# Viral attachment blocking chimera composed of DNA origami and nanobody inhibits Pseudorabies Virus infection *in vitro*

**DOI:** 10.1101/2023.02.13.528373

**Authors:** Swechchha Pradhan, Carter Swanson, Chloe Leff, Isadonna Tengganu, Melissa H. Bergeman, Ian B. Hogue, Rizal F. Hariadi

## Abstract

Antivirals are indispensable tools that can be targeted at viral domains directly or indirectly at cellular domains to obstruct viral infections and reduce pathogenicity. Despite their transformative use in healthcare, antivirals have been clinically approved to treat only 10 of the more than 200 known pathogenic human viruses. Additionally, many virus functions are intimately coupled with host cellular processes, which present challenges in antiviral development due to the limited number of clear targets per virus, necessitating an extensive insight into these molecular processes. Compounding this challenge, many viral pathogens have evolved to evade effective antivirals. We hypothesize that a Viral Attachment Blocking Chimera (VirABloC) composed of a viral binder and a bulky scaffold that sterically blocks interactions between a viral particle and a host cell may be suitable for the development of antivirals agnostic to the extravirion epitope that is being bound. We test this hypothesis by modifying a nanobody that specifically recognizes a non-essential epitope presented on the extra virion surface of Pseudorabies virus strain 486 with a 3-dimensional wireframe DNA origami structure ∼100 nm in diameter. The nanobody switches from having no inhibitory properties (tested up to 50 µM) to 4.2 ± 0.9 nM IC_50_ when conjugated with the DNA origami scaffold. Mechanistic studies support that inhibition is mediated by the non-covalent attachment of the DNA origami scaffold to the virus particle, which obstructs the attachment of the viruses onto host cells. These results support the potential of VirABloC as a generalizable approach to developing antivirals.

## Introduction

Beginning in the 1960s^1^, antiviral therapeutics have been developed to address circumstances where vaccines are not useful, such as targeting viruses for which vaccination is not effective (e.g., HIV or Herpes infections), not safe (e.g., the risk of antibody-dependent enhancement of dengue virus infection), the time to develop protective immunity is limited (e.g., during a viral outbreak), or for use in immunocompromised patients. For example, antivirals have revolutionized the treatment of HIV/AIDS, significantly reducing morbidity and mortality to such an extent that HIV is now considered a largely manageable chronic infection. ^2^ While the rate of drug discovery has increased substantially in recent decades, the timeline for clinical testing and authorization by regulatory authorities remains the major bottleneck. The current arsenal of antiviral treatments contains only about 118 FDA-licensed antiviral drugs that treat only ∼ 10 out of more than 200 human viral infectious diseases. ^3^ Even more concerning is the possibility that post-authorization, the lifetime of these drugs will be severely cut short due to the intractable problem of drug resistance. The large population size of viruses, their high mutation rates, and the often unchecked exposure of animal reservoirs to antiviral drugs work together to accelerate virus adaptation to resistance. ^4^

Conventional efforts to develop antivirals have focused on detailed structure-function relationships between a therapeutic and the targeted viral or host proteins. ^5–9^ One approach is to utilize the binding of antibodies to an essential epitope to competitively inhibit interactions with host cells. However, insufficient functional insight is available for many viruses to implement such a targeted development strategy. Compounding this challenge, the emergence of new viral strains and variants may render previously developed antivirals ineffective. For example, during the ongoing COVID-19 pandemic, none of the monoclonal antibody therapeutics developed to treat the earlier variants of SARS-CoV-2 remain effective against later variants. ^10^ Here we consider a generalizable approach for developing an antiviral, which does not require great mechanistic insight for a given viral pathogen and allows the use of virus binders that would not typically be considered useful for the development of therapeutics. Several methods have been described to develop chemical moieties that can specifically bind a virus of interest. For example, traditional hybridoma screening methods can discover monoclonal antibodies that specifically bind a virus of interest. While some monoclonal antibodies discovered this way may exhibit antiviral activities, most will not be suitable candidates for further antiviral development because they bind to viral epitopes that are non-essential for viral infection.

The development of proteolysis targeting chimeras (PROTACs), wherein chemical molecules can be conjugated to form a bifunctional chimeric molecule with a rationally designed function, has spurred much research interest. ^11^,^12^ By designing a drug-like molecule with different functional groups, new candidates have emerged for previously undruggable proteins. We hypothesize that by breaking down the design of viral inhibitors into two functions, (1) binding to the virus of interest and (2) blocking the attachment of that virus to a host cell, a chimeric molecule may be assembled rationally and may similarly aid in the development of new antiviral candidates. We term the approach applied in this report as a Viral Attachment Blocking Chimera or VirABloC (Figure 1).

**Fig. 1.**
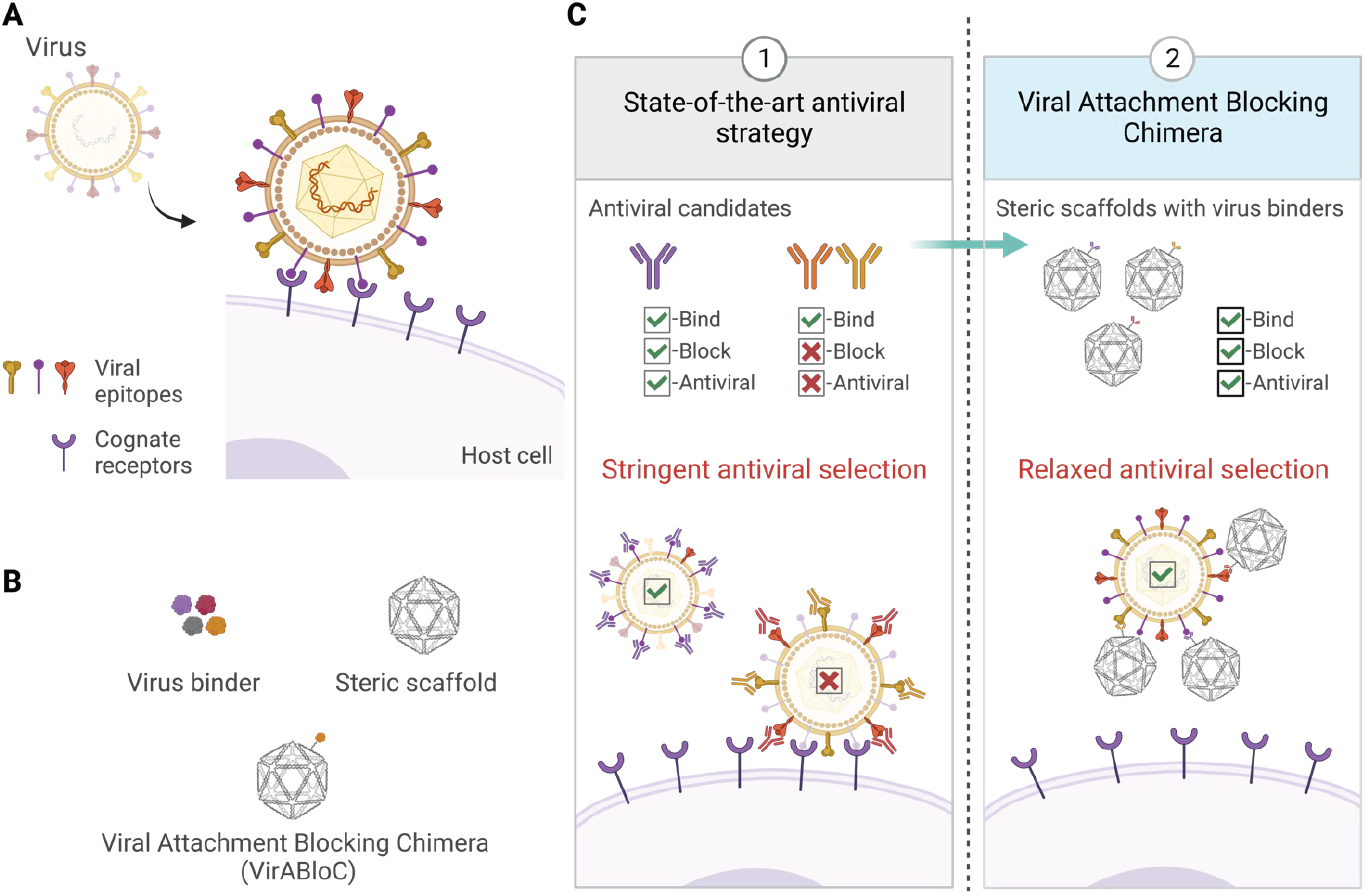
Design of Viral Attachment Blocking Chimera (VirABloC). (**A**) During virus infection, multivalent interactions between virus particles and host cell receptors mediate virus attachment and entry. (**B**) Two-component system of VirABloC comprising virus binding domains (binders) and the steric scaffold. (**C**) In the current state-of-the-art, only a subset of viral binders (e.g., neutralizing antibodies) block critical viral protein functions to have an antiviral effect (**1**). By functionalizing an otherwise less-ideal virus binder with a steric scaffold, low-affinity binders targeting non-essential viral epitopes can have antiviral effects (**2**).

To test our hypothesis, we first sought to identify a viral binder that targets a specific epitope but binds with relatively low affinity and has no intrinsic antiviral activity. We chose to use a nanobody that targets GFP, which is unlikely to cross-react with other native viral epitopes. We then chose to test Pseudorabies Virus (PRV; suid herpesvirus 1) as a model for human Herpes Simplex 1 and 2 (HSV-1 and -2) and because it is an economically important veterinary virus. These herpesviruses present a unique challenge for prevention and treatment because they establish lifelong latent infections, vaccine candidates against human HSV-1 and -2 have largely proven ineffective, and vaccination of livestock against PRV has been frustrated by vaccine escape mutants. The recombinant viral strain PRV 486 contains pHluorin, a variant of GFP, inserted into an extravirion loop of the non-essential envelope glycoprotein gM. ^13^ We chose this binder and viral strain system because gM-pHluorin is incorporated into virus particles yet is not essential for viral infectivity and replication. In addition, pHluorin contains mutations in the epitope bound by the anti-GFP nanobody, making it likely a relatively low affinity binder. Because of these qualities, the nanobody binder is not expected to have any antiviral activity on its own. Such a combination of attributes is intended to model a typical, less desirable epitope that might emerge from an uninformed screen of binders, which would not be expected to make an effective antiviral therapeutic on its own.

Next, we considered the scaffold composition. Recently, several groups have reported DNA origami with various geometries multivalently conjugated with antiviral binders. ^14–16^ DNA origami is an ideal approach due to the ease of conjugating biomolecules, the control of the number and orientation of biomolecules, and its compatibility with biological models. ^17–22^ We chose a DNA origami wireframe structure that assembles into an icosahedron shape termed ‘snub cube,’ with a diameter of ∼ 60 nm ^23^, because of its large volume, low density, and spatial separation of potential conjugation sites. In our study, the snub cube was designed to have either 1, 12, or 60 single-stranded, outwardly rectifying DNA ‘handles’ evenly dispersed across the surface, where the anti-GFP nanobody conjugated to a complementary single-stranded DNA oligonucleotide binds.

In this study, we report that scaffolding the anti-GFP nanobody with the DNA origami snub cube switches it from inactive to a potent antiviral in an *in vitro* model of viral infection. We find that it inhibits infection by blocking PRV attachment to host cells. Additionally, multivalent conjugation of anti-GFP nanobody to the snub cube scaffold can be used to enhance PRV 486 binding and IC_50_ from low µM to low nM. These findings suggest modifying viral binding molecules with a bulky scaffold may be a generalizable approach to developing antivirals by blocking viral attachment to host cells.

## Results

### Specific binding of nbGFP binds to PRV 486

We first set out to characterize the binding of anti-GFP nanobody (nbGFP) to PRV 486. Previous studies have reported that the binding affinity of nbGFP to EGFP to be *K*_*D*_ < 2 nM. ^24–26^ In this study, we sought a molecular epitope that retains the specificity of this interaction but has a lower binding affinity. Superecliptic pHluorin is a fluorescent protein derived from GFP with a cluster of mutations that makes its bimodal excitation (395 and 475 nm) and emission (509 nm) sensitive to changes in the pH. 27 A crystal structure of EGFP and nbGFP has been solved, revealing the precise binding interface between these two proteins. Amino acid residues 147, 204, and 206 are found at the epitope that binds to nbGFP and differs between pHluorin and EGFP. Despite these modifications, nbGFP has been successfully used to detect pHluorin-tagged proteins ^28^, and PRV 486 viruses have been detected using anti-GFP antibodies for immunofluorescence (IF) studies. ^13^ We anticipate that nbGFP will specifically bind to the pHluorin domain on PRV 486 but with a substantially lower affinity than EGFP.

We used a fluorescence microscopy-based assay to image the immunocapture of either monomeric EGFP or PRV 486 by either nbGFP or an anti-GFP monoclonal antibody immobilized on the surface of the slide (Figure 2A). As a positive control, EGFP protein was efficiently bound by both the anti-GFP monoclonal antibody and nbGFP (Figure 2B). PRV 486 particles, which display the pHluorin domain on their virion surface, were also captured by the monoclonal antibody and nbGFP (Figure 2B). These characteristics indicate that the nbGFP-PRV 486 system is a suitable virus-binder pair for our study.

**Fig. 2.**
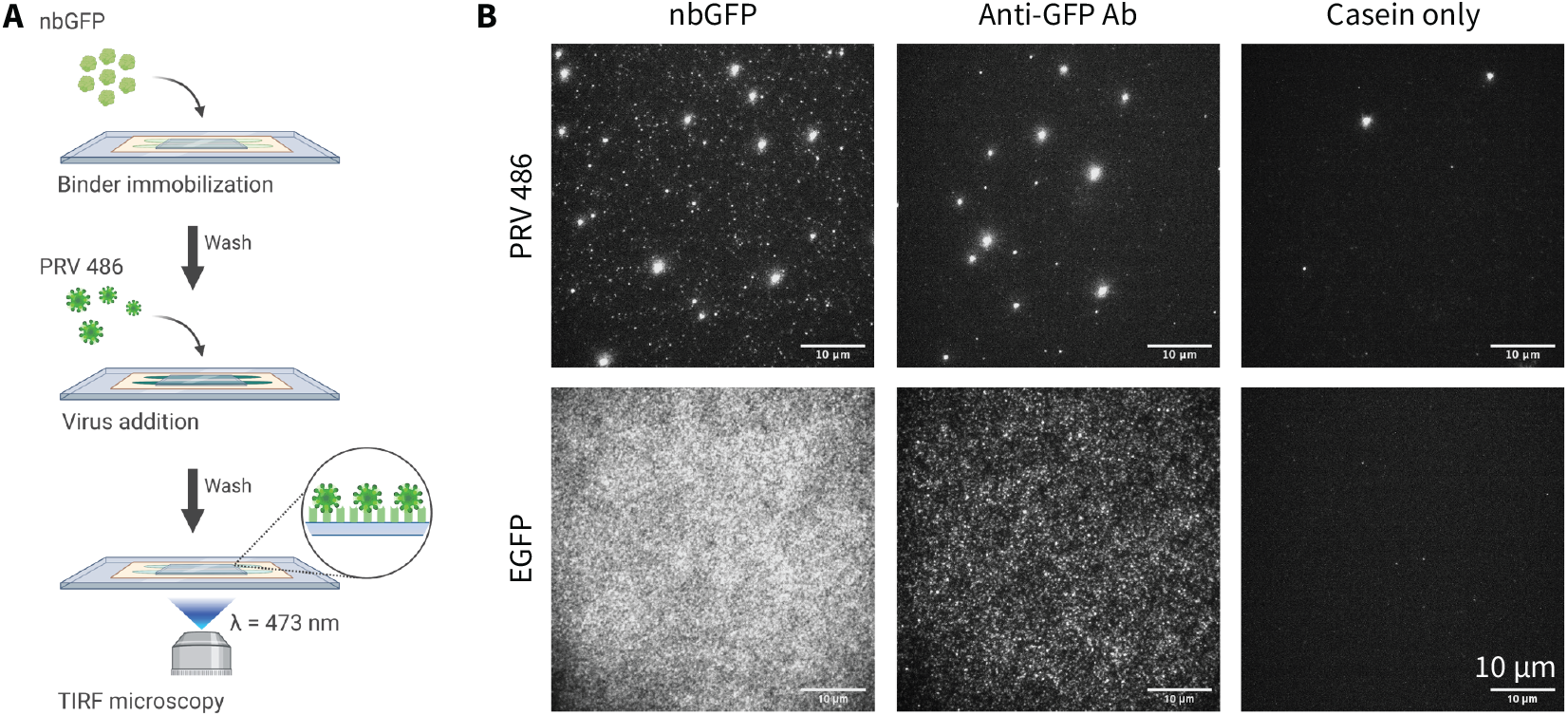
nbGFP binds to PRV 486. (**A**) Schematic illustrating the capture assay. The nbGFP binder was immobilized on the glass surface. PRV 486 expresses pHluorin on the outside of the viral envelope. The capture of PRV 486 particles on the glass surface was measured by TIRF microscopy. (**B**) Representative TIRF microscopy images showing binding between nbGFP and PRV 486. Anti-GFP antibody is used as the positive control for nbGFP, and EGFP is used as the positive control for PRV 486. Casein (non-specific blocking agent) is used as a negative control for nbGFP. The scale bar is 10 µm.

### Design, synthesis, and characterization of DNA origami-based multivalent binder

The choice of scaffold is critical for designing multivalent binder/inhibitor systems. In this study, we chose a 3D DNA origami scaffold to control the spatial distribution of virus binders with defined nanometer-level precision and to minimize batch-to-batch variations of the assembled nanostructures (Figure 3A). Previous studies have shown that the size, shape, and complexity of DNA nanostructures affect their stability, pharmacokinetics, and interactions at biological interfaces *in vitro* and in vivo. ^29–36^ Like other herpesviruses, PRV particles are about 200 nm in diameter. Studies demonstrating a size-dependent effect on multivalent interactions have shown a higher degree of viral inhibition with inhibitor sizes slightly smaller or similar to the virus particle. ^37^,^38^ Increasing the size of the DNA origami scaffold to match the size of the virus may increase the antiviral efficacy of the multivalent system and provide a higher surface area to increase the valency without clustering of ligands.^37^ However, a larger DNA origami nanostructure will likely compromise its immunotolerance and necessitate a higher Mg^2+^ ion concentration to maintain its structural integrity, which is not typical in physiological environments. 29,30 To rationally design the 3D scaffold, considering the size range of virus particles, maintaining the structural integrity of the assembled multivalent system, and maintaining a low surface area to volume ratio, we opted for wireframe DNA origami scaffolds, 50 − 100 nm diameter size.

**Fig. 3.**
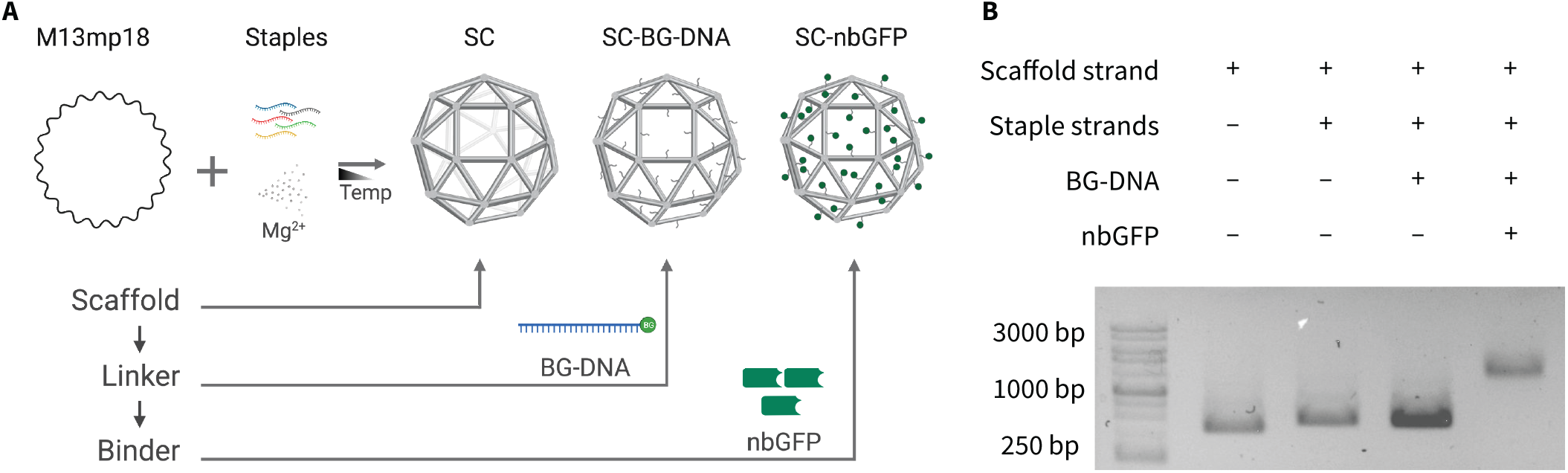
Design and synthesis of DNA origami and nanobody chimera. (**A**) Schematic showing the stepwise formation of SC-nbGFP using DNA origami technology. The technique involves folding the M13mp18 ssDNA using short staple strands in the presence of cations such as Mg^2+^. The 3D rod structure of the snub cube (SC) with 24 vertices and 60 edges is shown on the right. The second step is the incorporation of the BG-DNA bifunctional linkers onto the SC (front face only), followed by the multivalent conjugation of nbGFP virus binders. (**B**) Agarose gel electrophoresis showing the successive formation of SC60H, SC60H-BG-DNA, and SC60H-nbGFP. The unfolded M13mp18 strand is used as a reference.

To this end, we adapted the snub cube (SC), a 3D wireframe DNA origami, ∼ 60 nm diameter size, with 60 edges, 24 vertices, and 38 faces, including 6 squares and 32 equilateral triangles (Figure 3B and Figure S1).^23^ We modified the original SC design using Tiamat ^39^ to incorporate 20-nt single-stranded DNA (ssDNA) overhangs in the middle of its edges. To functionalize SC with virus binders, we site-specifically conjugated one or more copies of the SNAP-tag nbGFP per SC (one binder on one edge of the SC) via benzyl guanine (BG) linkers (Figure 3C). Using this approach, we could vary the copy number of virus binders in the SC scaffold from 1 to 60, assuming a 1:1 stoichiometric ratio of binding between the nbGFP and SC edges. To span the range between 1 and 60, we constructed SC-nbGFP of three valences, including SC1H-nbGFP, SC12H-nbGFP, and SC60H-nbGFP. We maintained a 12.5 mM Mg^2+^ concentration throughout all preparation and purification steps to retain the structural integrity of the assembled nanostructures. We characterized the formation of SC, SC-BG-DNA, and SC-nbGFP after each step using agarose gel electrophoresis. The formation of distinct bands and the reduced electrophoretic mobility of these bands were consistent with their relatively increasing molecular weights, confirming the correct formation of nanostructures after each synthesis step (Figure 3D and Figure S2). The final SC-nbGFP exhibited the lowest mobility in gel electrophoresis, followed by SC-BG-DNA intermediate nanostructures with the complementary ssDNA attached to the BG linker, and finally by the SC constructs with 60 ssDNA overhangs corresponding to their valency. In addition, we characterized SC60H-nbGFP using dynamic light scattering (DLS) and found a majority peak at ∼ 100 nm in diameter (Figure S1).

### Multivalent conjugation of nbGFP to SC enhances virus binding

Since multivalency can be leveraged to offset weak binding, we expected the multivalent SC-nbGFP nanostructures to have comparatively stronger binding affinities than monomeric nbGFP for PRV 486. To address the functionality of the multivalent system, we first analyzed the impact of SC-nbGFP valency on virus binding using semiquantitative ELISA assays (Figure 4). We coated 96-well ELISA plates with PRV 486 and incubated them with monomeric nbGFP and multivalent SC-nbGFP nanostructures of different valences. We quantified the extent of binding of nbGFP and SCnbGFP nanostructures using an orthogonal nbGFP-specific reporter antibody.

**Fig. 4.**
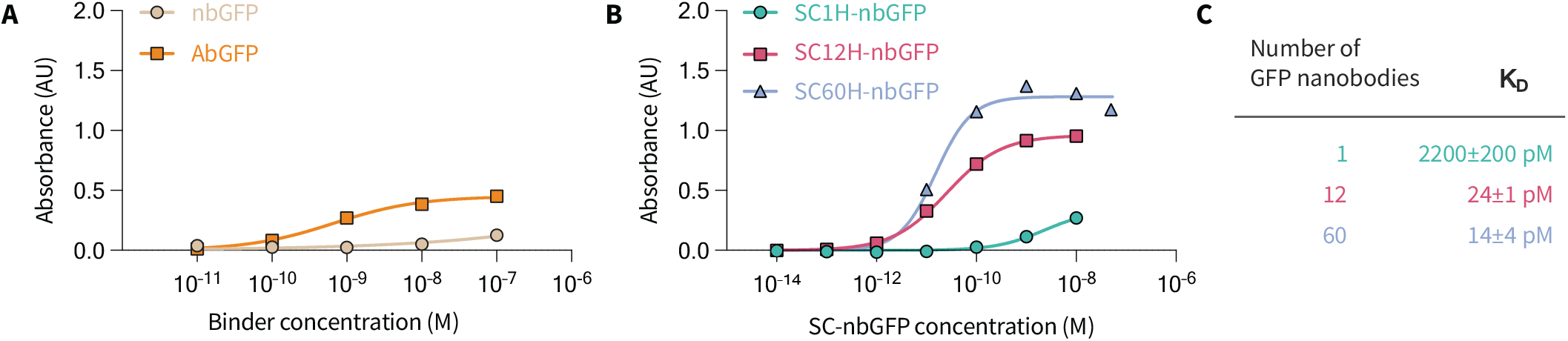
Multivalent presentation of nbGFP on DNA origami scaffold increases viral binding. (**A**) ELISA binding curves of PRV 486 to nbGFP compared to a monoclonal antibody. (**B**) ELISA binding curves of PRV 486 to SC-nbGFP of different valences (*R*^2^ >0.95). Absorbance values resulting from non-specific interactions without the virus or EGFP were subtracted. The binding affinities of SC1H-nbGFP, SC12H-nbGFP, and SC60H-nbGFP calculated by non-linear regression are listed in (**C**).

The monomeric nbGFP, monovalent SC1H-nbGFP, and multivalent SC12H-nbGFP and SC60H-nbGFP showed concentration-dependent changes in signal intensities, suggesting specific binding activity (Figure 4). As expected, nbGFP exhibited a relatively low affinity for the pHluorin moiety on PRV 486, most likely due to the sequence differences between pHluorin and EGFP. The low binding of nbGFP at concentrations <10 nM may also be attributed to inefficient passivation of the virus surfaces to prevent nonspecific interactions (Figure 4A). However, the multivalent interactions of SC12H-nbGFP and SC60H-nbGFP increased the overall binding affinity, consistent with previously published studies using multivalent 2D and 3D DNA nanostructures for virus sensing and inhibition. ^16,40^ The non-linear fits are the specific binding with Hill slope *h* given by

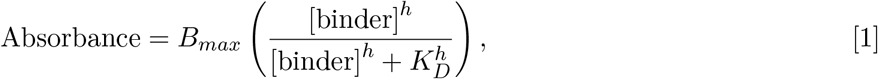

where *h* is the Hill slope, *B*_*max*_ is the effective maximum specific binding, and K_D_ is the binder concentration needed to achieve a half-maximum binding at equilibrium. The relative K_D_’s showed a 90-fold increase in the binding affinity of SC12H-nbGFP (24±1 pM) and a 150-fold increase in the binding affinity of SC60H-nbGFP (14±4 pM) compared to that of monovalent SC1H-nbGFP (2200±200 pM; Figure 4B) and GFP antibody (670±10 pM; Figure 4A). These data support our conclusion that nbGFP targets the pHluorin-tagged gM domains of the PRV 486, and multivalent conjugation of nbGFP onto SC enhanced the binding strength of the SC-nbGFP complex with PRV 486.

### Multivalent SC-nbGFP inhibits PRV 486 infectivity *in vitro*

We next addressed our hypothesis that the attachment of SC-nbGFP onto the viral particle will inhibit viral infection. To investigate this hypothesis, we used dose-response analysis using plaque reduction assays *in vitro* (Figure 5A). We first evaluated whether nbGFP alone, SC alone, or nbGFP and SC not covalently conjugated could disrupt plaque formation. We found that nbGFP alone, up to concentrations of 50 µM, has no impact on plaque formation (Figure 5B). Similarly, SC alone, or SC and unconjugated nbGFP, had no impact on plaque formation up to 50 nM of SC and 15 µM of nbGFP (Figure 5C). The fitted curves for the infection rates *I*’s in Figures 5, 6, S3, and S4 are

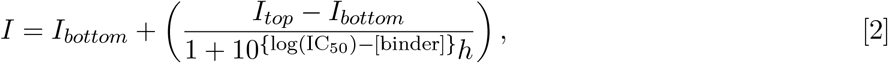

where *I*_*top*_ and *I*_*bottom*_ are fitting parameters that correspond to the plateaus of infectivity *I, h* is the Hill slope, and IC_50_ is the concentration of binder that gives infectivity between *I*_*top*_ and *I*_*bottom*_. In contrast, SC60H-nbGFP demonstrated dose-dependent inhibition of PRV 486, with an estimated half-maximal inhibitory concentration (IC_50_) of 4.2±0.9 nM (Figure 5D). The measured IC_50_ is within a factor of 2 of K_D_ between nbGFP and PRV. These results support our central hypothesis that the attachment of a DNA origami constructs to a non-essential viral epitope binder can disrupt viral infection. Furthermore, these results show that both specific (albeit low affinity) binding, mediated by nbGFP, and multivalency/steric size, conferred by SC, are required for effective inhibition.

**Fig. 5.**
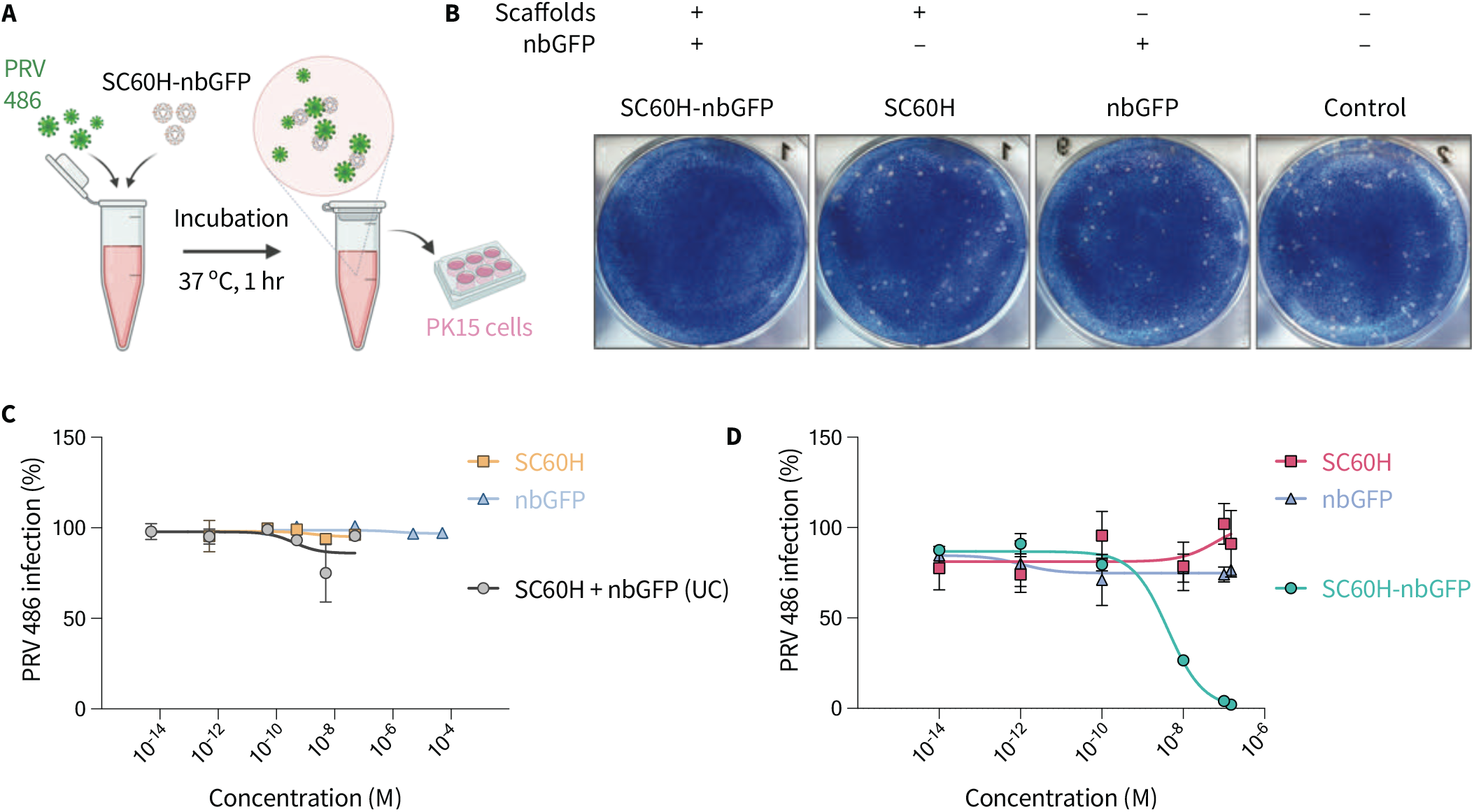
SC60-nbGFP inhibits PRV infectivity *in vitro*. (**A**) Schematic illustrating infectivity assays. (**B**) Representative plaque assays corresponding to the maximum concentration of SC60-nbGFP, SC60, and nbGFP at 150 nM. Untreated PRV 486 is shown for reference. (**C**) Dose-dependent, plaque-reducing inhibition curves for control groups including nbGFP (monomeric binder only), SC60H (scaffold only), and unconjugated (UC) SC60H with 5X molar excess of nbGFP. Error bars represent mean ± SD, N=3 biologically independent experiments. (**D**) Dose-dependent, plaque-reducing inhibition curves for the SC60-nbGFP, SC60, and nbGFP. Error bars represent mean±SD, N=2 biologically independent experiments.

**Fig. 6.**
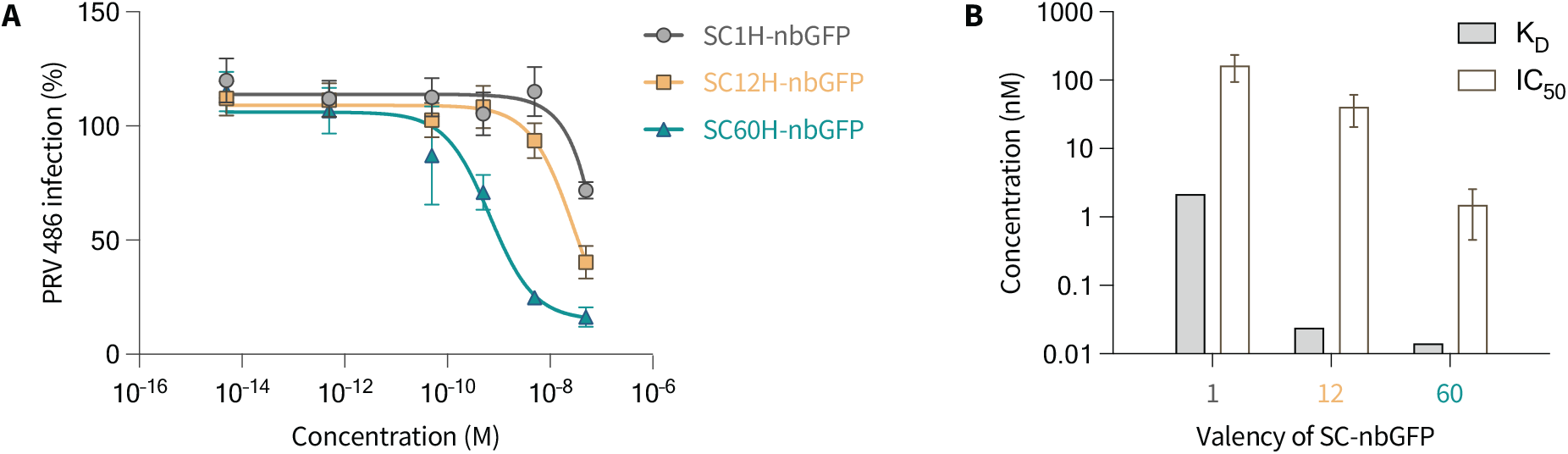
SC-nbGFP inhibition correlates with binder valency and affinity. (**A**) Dose-dependent, plaque-reduction inhibition curves for PRV 486 incubated with SC-nbGFP nanostructures of valency, n = 1, 12, and 60. Data are presented as mean ± SD, N = 3 biologically independent experiments. (**B**) Comparison of binding affinity (K_D_) and inhibition efficiency (IC_50_) of SC-nbGFP for different valences.

### Multivalency enhances IC_50_, but monovalency is sufficient for antiviral effects

We next sought to understand the mechanism of viral inhibition. One hypothesis is that inhibition is a function of multivalency. Such a conclusion was reported in recent studies using DNA scaffolds in which multivalent interactions and spatial pattern matching were used to offset weak binding affinities of virus-inhibiting aptamers and achieve efficient viral inhibition with enhanced IC_50_ values. ^14,16,41^ In these studies, the topological patterning of virus binders shielded virus particles from interacting with cell surface receptors. Another study reported that a multivalent 3D DNA nanostructure caused aggregation of virus particles, reducing the effective concentration of infectious viruses.

To investigate the impact of valency, we carried out dose-response analyses of PRV 486 inhibition with either SC1H-nbGFP, SC12H-nbGFP, or SC60H-nbGFP, and measured viral infection by flow cytometry 48 hours post-infection. Virus inhibition was negligible with nbGFP or SC scaffolds alone, and unconjugated nbGFP + SC (Figure S3). On the contrary, when conjugated, the SC-nbGFP constructs showed dose-dependent inhibition of the virus (Figure 6B). The inhibition efficiency increased monotonically with the rising valency of SC-nbGFP, with IC_50_ values of 150 nM for SC1H-nbGFP, 31± 6 nM for SC12H-nbGFP, and 0.67 ± 0.02 nM for SC60H-nbGFP. The IC_50_ of SC60H-nbGFP measured using flow cytometry in this experiment is similar to that previously obtained using the plaque reduction assay in Figure 5 (4.2 ± 0.9 nM). Our findings indicate a positive correlation between K_D_ and IC_50_ of the SC-nbGFP nanostructures (Figure 6B). This is consistent with numerous other studies with multivalent inhibitors^42–44^ and is likely explained by the increased avidity resulting from the multiple nbGFPs per SC, leading to enhanced binding and higher inhibitory potency. However, there is also an important distinction: the observed inhibition with monovalent SC1H-nbGFP and the absence of inhibition with unconjugated monomeric nbGFP suggest that the platform does not simply function by increasing avidity. Instead, inhibitory effects require the presence of an SC construct, even with low-affinity monovalent binding.

To determine whether multivalent SC-nbGFP causes cross-linking and aggregation of virus particles, we analyzed the particle size distribution of viruses incubated with SC60H-nbGFP using a nanoparticle analyzer, which measures the diffusion coefficient of individual particles to estimate their effective hydrodynamic size (Figure S4). PRV has previously been reported to have a diameter of ∼ 200 nm. ^45^ Using the nanoparticle analyzer, we observe that our stock PRV 486 exists as a single monodisperse peak of ∼ 200 nm. In comparison, all the other groups also showed a uniform size distribution of particles centered around ∼ 200 nm, with negligible formation of higher-order aggregates. The lack of observed aggregation is expected since in this work, the viral concentration was typically chosen to be on the order of 1 pM. A minuscule number of virus particles leads to a low collision frequency between viral particles and slow nucleation and polymerization kinetics of viral particles. Interestingly, while Dynamic Light Scattering (DLS) showed that SC formed with expected size profiles (Figure S1), binding of SC-nbGFP to virus particles did not appear to increase their hydrodynamic size as measured by the NTA.

Our studies of valency found that only one nbGFP per SC was sufficient to produce antiviral effects. This suggests that (a) not all binding sites on the virus are saturated since the size of SC would likely sterically block access from other SC-nbGFP and (b) crosslinking interactions between two or more viral particles mediated by SC-nbGFP are not a critical feature of the inhibition mechanism. These pieces of data provide strong evidence that SC-nbGFP does not function through the aggregation of PRV 486.

### SC-nbGFP obstructs viral attachment to host cell surfaces

Previous work on PRV has shown that virus particles initiate infection by adhering to target cells, then internalizing into the cytoplasm primarily via membrane fusion at the plasma membrane or via the endocytic pathway.^45–47^ In either case, viral capsids are internalized into the cells and transported to the nucleus. Since we targeted pHluorin domains located on the outside of the PRV 486 virion envelope using SC-nbGFP nanostructures, we anticipated that the multivalent interactions would obstruct the entry events of PRV particles, resulting in reduced attachment, reduced internalization, or both.

To assess whether SC-nbGFP blocks PRV 486 surface attachment to host cells, we performed time-lapse live-cell fluorescence imaging to track the early events of virus infection. To this end, we used PRV 483, a different recombinant strain expressing gM-pHluorin on the virus envelope, and a red fluorescent mRFP-VP26 virus capsid tag. At neutral pH, PRV 483 particles exhibit colocalized green and red fluorescence in the extracellular space but shed their virion envelope and exhibit red-only fluorescence after internalization (Figure 7A). Using PRV 483, we monitored colocalized green and red extracellular puncta versus internalized red-only puncta at 0, 30, and 60 mins post-infection. To be able to see a sufficient number of virus particles in this experiment, we used a multiplicity of infection (MOI) of 10^4^ infectious units per cell. To quantify attachment efficiency, we counted the total GFP puncta per unit area, comparing two experimental conditions: PRV 483 pre-incubated with SC-nbGFP, and PRV 483 pre-incubated with SC alone. When treated with SC-nbGFP, we observed significantly less green puncta (pHluorin) per unit area compared to the control condition. The untreated viruses showed a 2-to 3-fold greater accumulation of pHluorin puncta per unit area over time (Figure 7B). However, SC-nbGFP did not decrease the number of internalization events per attached particle compared to the control (Figure S7). These data support our conclusion that SC-nbGFP obstructs the attachment of virus particles onto the host cell. Based on this observation, we termed this approach Viral Attachment Blocking Chimera (VirABloC). The observed inhibition is most likely achieved by sterically blocking attachment factor interactions between viral envelope glycoproteins and cellular heparan sulfate proteoglycans.

**Fig. 7.**
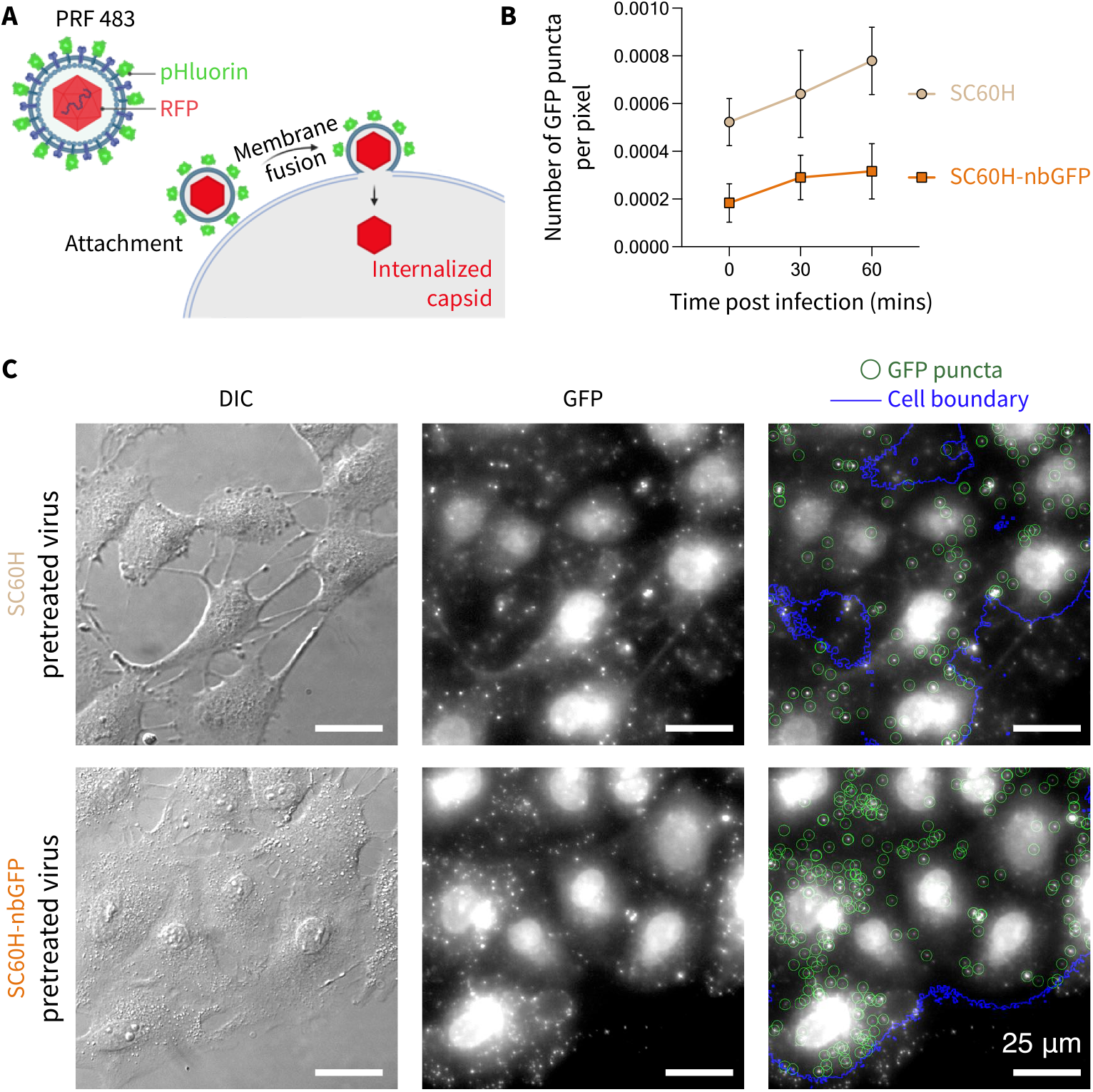
SC-nbGFP blocks viral attachment to host cells. (**A**) Schematic showing the early entry events of PRV 483 comprising attachment followed by membrane fusion and internalization of virus capsids inside the cytoplasm. (**B**) Attachment of virus particles pretreated with SC60H-nbGFP versus SC60H (control). Error bars are mean ± S.D. (**C**) Representative DIC and widefield fluorescence microscopy images of PK15 cells infected with PRV 486 pretreated with either SC60H-nbGFP (top row) or SC60H (bottom row) at 30 mins post-infection. Representative images at t = 0 and 60 mins can be found in Figure S7. A custom code in Mathematica (available at https://github.com/rhariadi/virabloc) marks the cell boundary and PRV 486 viral particles with blue lines and green circles, respectively.

## Discussion

In this study, we propose a novel antiviral platform. We demonstrate that a chimeric and bi-functional complex, composed of protein that specifically binds to a viral epitope, conjugated multivalently onto a steric-blocking scaffold composed of DNA origami, can be leveraged to expand the repertoire of antiviral targets to include non-essential viral domains. Our multivalent scaffold conjugated with nanobody showed an IC_50_ for PRV 486 in the low nanomolar range (3 nM), consistent with recent studies using multivalent DNA-based 2D and 3D scaffolds for antiviral applications. ^14–16^ Our findings suggest a more nuanced and active role of the scaffold than being a mere inert backbone for the spatial presentation of virus binders. In our study, the scaffold was targeted to a non-essential viral epitope and indirectly interfered with viral attachment, without needing to specifically target the viral factors responsible for attachment. It also suggests that this strategy will likely work with low-affinity binders by leveraging multivalency to offset weak binding, given that the binders are highly selective for their target viral epitopes. In addition, the avidity resulting from multivalency can be instrumental in further enhancing the platform’s inhibition potency. The assembled nanostructures demonstrated negligible cytotoxicity (<10 nM) and sufficient stability, further supporting their therapeutic potential (Supplementary Figure 4 and Supplementary Figure 6).

Since no antivirals are available for PRV, we could not directly compare our system’s efficacy. However, a recent paper reported an IC_50_ of 15.2–31.6 µg/mL (100− 600 nM) with monoclonal antibodies developed against gB of different PRV strains. ^48^ More importantly, in contrast to all the previous multivalent inhibitor designs that rely on binders targeting essential viral proteins, our multivalent system targeted the PRV gM. This glycoprotein is conserved throughout the Herpesviridae family but is not known to actively participate in virus entry processes. ^49^ To our knowledge, the therapeutic potential of non-essential viral targets has not been explored. For instance, in a recent report describing a system called “DNA Star”, DNA aptamer binders targeted the envelope protein domain III (ED3) clusters of the Dengue virus that interacts with the primary and secondary cell surface receptors.^50,51^ Similarly, the aptamer binders in the recent “DNA net” study targeted the receptor binding domains of the trimeric spike proteins of SARS-CoV-2.^16^ Similarly, numerous studies on influenza viruses have utilized proteins, peptides, and aptamers targeting the hemagglutinin (HA) epitopes that bind to the cell surface receptors and mediate the virus entry processes.^6,44,52,53^ As expected, by targeting these essential domains, the monovalent binders demonstrated antiviral activity as well, although poorly. The monomeric aptamers in the DNA star and the DNA net papers showed IC_50_ of ∼ 10 µM and ∼ 15 µM, respectively. In comparison, the multivalent inhibitors enhanced the IC_50_ values by 3 orders of magnitude. The enhancement was mainly attributed to their design strategy to display the binders matching the spatial pattern of virus epitopes with nanometer-scale precision, which relies on precise structural knowledge of the target virus. DNA nanostructures are an ideal chemistry for the rational design of multivalent inhibitors, a persistent challenge for conventional polymeric and inorganic scaffolds. In contrast to previous studies, the monomeric nbGFP binders used in our study failed to inhibit PRV 486 viruses even at a concentration of 50 µM (the highest concentration we could test), while the multivalent SC-nbGFP demonstrated an IC_50_ ∼ 3 nM. We demonstrated that a low-affinity binder, targeting a non-essential viral domain, which does not have any detectable antiviral activity can be switched to a highly potent inhibitor by conjugating it to a 3D DNA origami scaffold.

A possible explanation for this could lie in the spatial proximity of gM to other glycoproteins, such as gB, gC, gD, and gH/gL, which are responsible for PRV attachment and membrane fusion 44. Although the spatial distribution of PRV envelope glycoproteins has not been resolved yet, studies of other herpesviruses have shown that glycoproteins gB, gC, and gH/gL are evenly distributed on the virus envelope, but their distribution may reorganize during binding and membrane fusion. 54 In our study, the VirABloCs could sterically block the interactions of entry-associated glycoproteins with the cell surface receptors. Based on our findings, we argue that steric shielding can effectively modulate viral infection. This opens the prospect of expanding viral inhibitors to make use of non-essential viral epitopes if they are accessible on the extravirion surface and expressed in sufficient copy numbers. Our strategy can be implemented with other pathogens to modulate infectivity with negligible impact on host cells. This strategy can also be instrumental in targeting viral pathogens that are novel, emerging, and understudied. Unlike rational design strategies to match spatial epitope patterns, which rely on extensive knowledge of the molecular nature of host-pathogen interactions, our strategy can be implemented with limited information on the spatial distribution of virus epitopes and with non-essential targets. The SC scaffold can be adapted to include pre-characterized virus binders such as virus-specific peptides, aptamers, nanobodies, and antibodies, with weak to moderate binding affinities but great pharmacodynamic and pharmacokinetic profiles, to enhance virus binding avidity and inhibition efficiency.

We report a modest binding affinity of DNA origami scaffolds with 1 nbGFP to PRV 486 (2200 ± 200 pM). However, upon increasing the valency of SC-nbGFP, the relative *K*_*D*_ dropped to 24 ± 1 pM for SC12H-nbGFP and 14 ± 4 pM for SC60H-nbGFP. With SC1H-nbGFP, since the range of concentrations we tested fell below the saturation point on the ELISA binding curve, we could not compare its binding affinity with the other groups. However, the avidities of other groups showed an enhancement in virus binding by three orders of magnitude compared to the monomeric nbGFP (Figure 3B). This enhancement of binding affinity upon increasing the valency of SC-nbGFP is consistent with previous studies using multivalent frameworks for virus binding. ^43,44,55,56^ Because DNA origami is a versatile framework for modulating conjugation sites, future work may explore the conjugation of binders targeting two or more distinct epitopes on the same viral target. Tri-valent inhibitors, where each binder recognizes a distinct viral epitope, have recently been reported to suppress antiviral resistance in SARS-CoV-2. ^9^ Using DNA origami, these studies could be further developed wherein the diversity of viral epitopes and binding avidity can be balanced in a single inhibitor composition. Future experiments could explore a polyclonal antibody mixture isolated from serum conjugated to a DNA origami scaffold that may expand the heterogeneity of viral epitopes targeted in the absence of structural insight into the viral target.^57^ While nanobodies and DNA origami served as valuable moieties in testing the VirABloC approach, future efforts in developing such viral inhibitors may not be constrained by these chemical types. Much work is needed to assess the generalizability of our findings, in particular, its capacity to function *in vivo* and for other viruses; however, our work demonstrates the feasibility of this approach.

## Acknowledgements

The authors thank Dr. C. Diehnelt for his assistance with the initial phase of the project. We thank the S. Sivaramakrishnan laboratory for their gift of the GFP nanobody plasmid, A. Varsani for helpful discussions, H. Glenn and T. Mukherjee for their assistance with confocal imaging, and T. Mukherjee and T. Taniguchi for their assistance with image processing and analysis.

## Correspondence

All correspondence and requests for materials should be addressed to R.F. Hariadi.

## Supporting Information

### S1. Supporting Figures

**Fig. S1.**
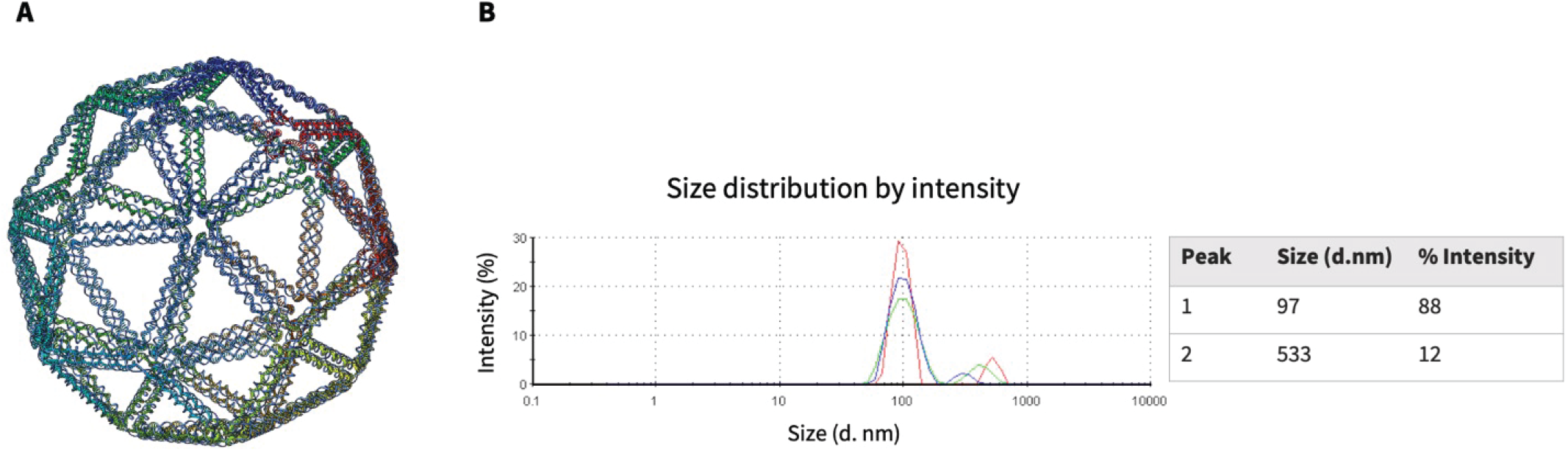
Characterization of SC and SC60H. (**A**) 3D model of the DNA origami snub cube. The model was generated using DAEDALUS. (**B**) Intensity particle size distribution (IPSD) of SC60H, analyzed using Dynamic Light Scattering, shows a majority peak at 97 nm diameter with no significant aggregation in the forms of trimers, tetramers, and higher order polymers. The 3D illustration in panel (**A**) is courtesy of Tohma Taniguchi.

**Fig. S2.**
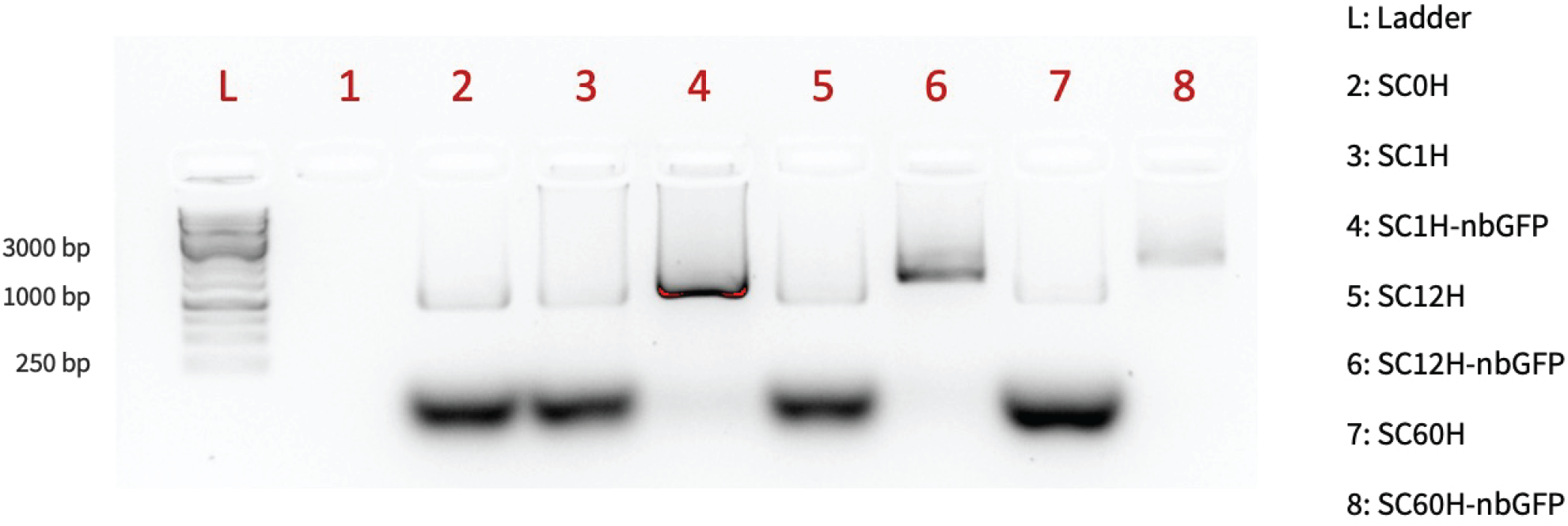
Characterization of SC-nbGFP constructs. (**A**) Agarose gel electrophoresis shows a shift in mobility due to the formation of SC1H-nbGFP, SC12H-nbGFP, and SC60H-nbGFP.

**Fig. S3.**
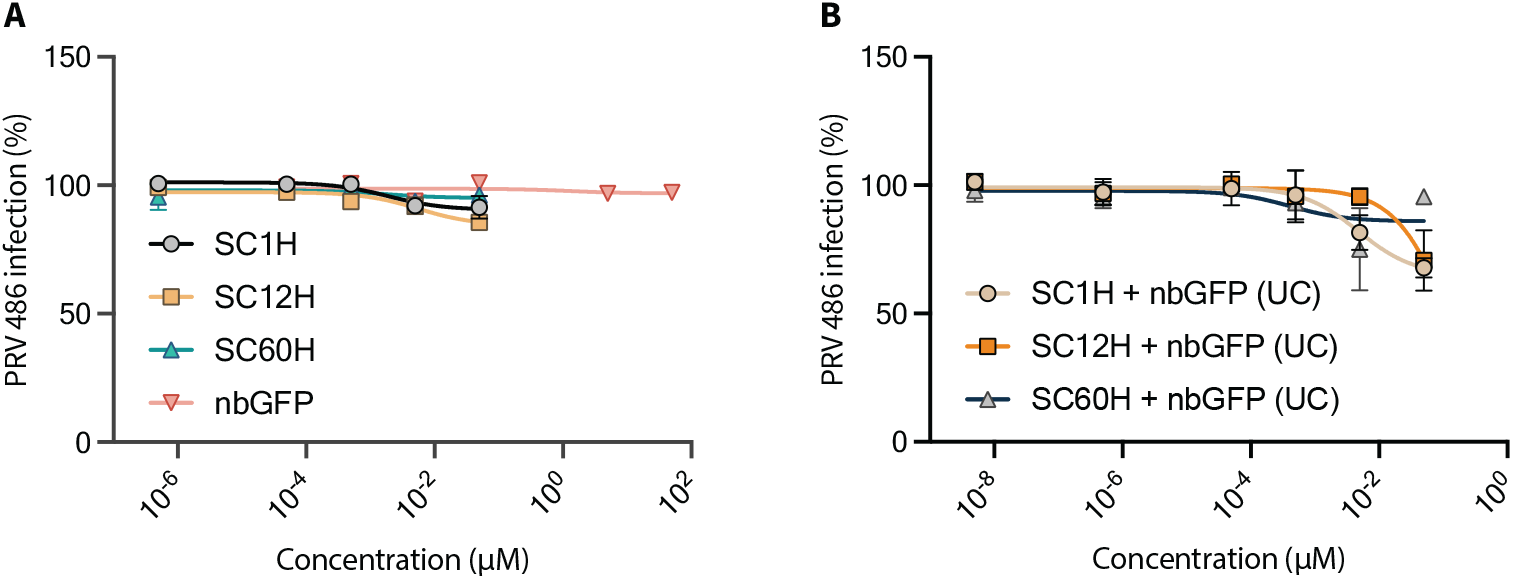
(**A**) Dose-dependent plaque-reduction inhibition curves for PRV 486 incubated with only SC scaffolds and monomeric nbGFP. (**B**) Dose-dependent plaque-reduction inhibition curves for PRV 486 incubated with unconjugated (UC) SC and nbGFP. Data are presented as mean ± S.E.M., N = 3 biologically independent experiments.

**Fig. S4.**
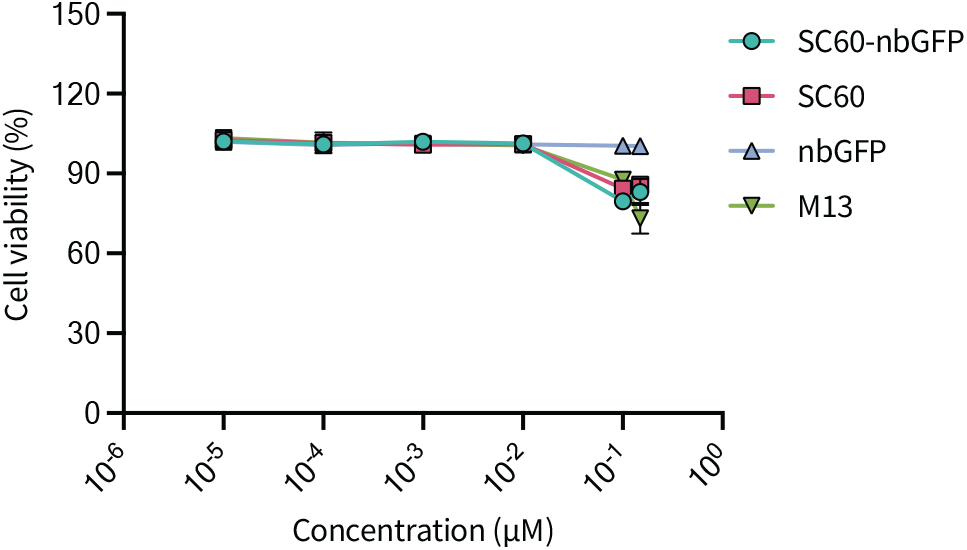
In vitro cytotoxicity assay. Cell viability after treatment with SC60-nbGFP for 24h. Data are presented as mean±S.E.M., N = 2 biologically independent experiments.

**Fig. S5.**
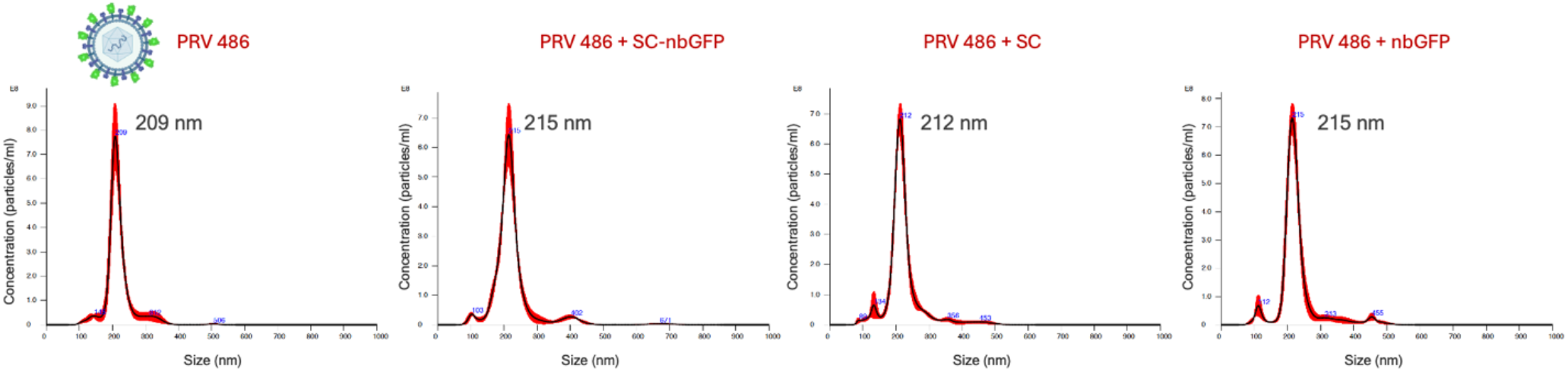
Size and aggregation characterization of SC-nbGFP and virus complexes. Nanoparticle tracking analyses (NanoSight) showing the particle size distributions of PRV 486 particles incubated with SC-nbGFP and components. The uniform size distributions indicate the absence of virus aggregation. For reference, PRV is ∼ 200 nm, and SC-nbGFP is ∼ 100 nm.

**Fig. S6.**
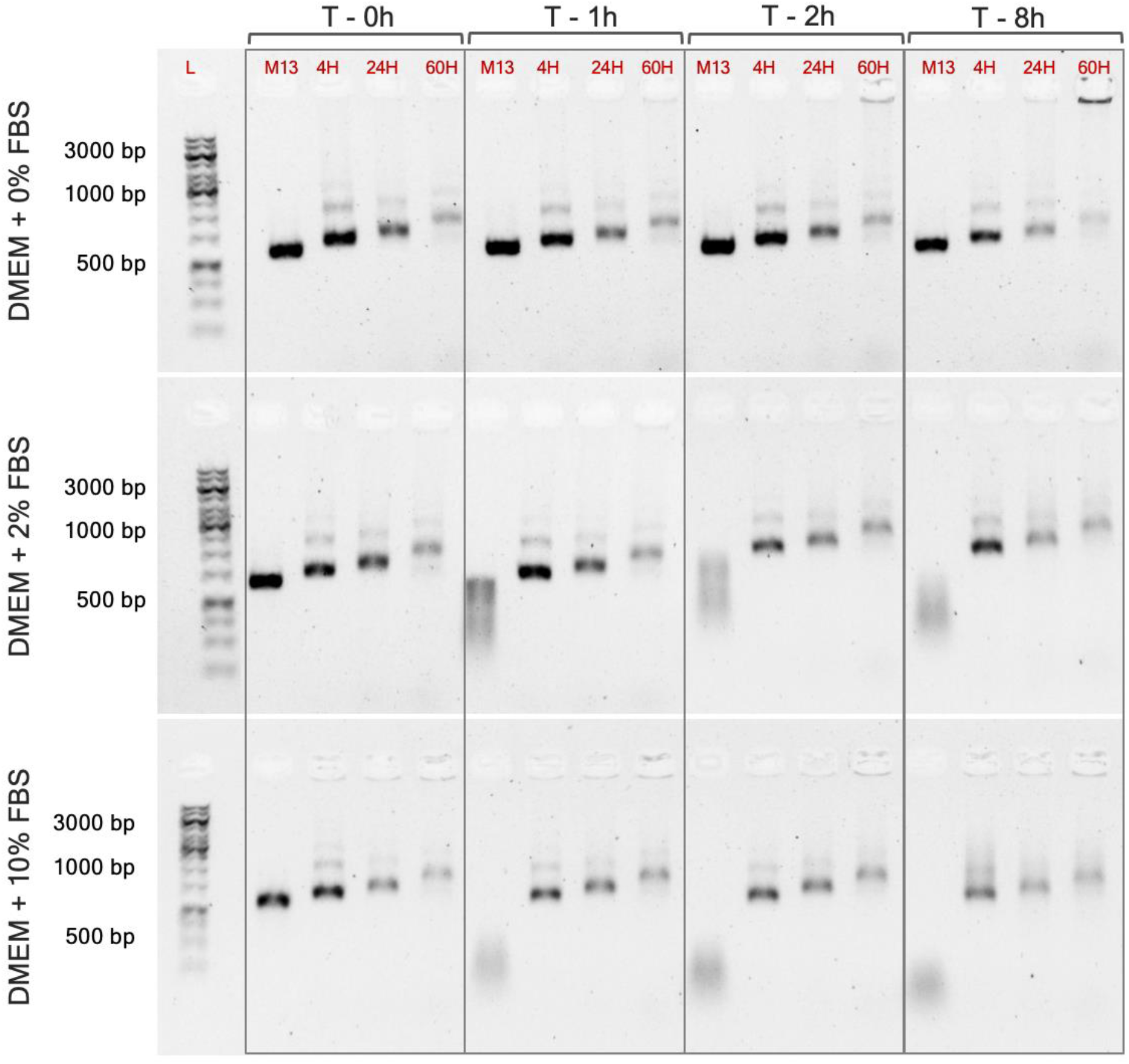
SC-nbGFP nanostructures are stable in serum for at least 8 hours. Agarose gel electrophoresis analysis of SC-nbGFP nanostructures incubated in DMEM supplemented with FBS (0%, 2%, and 10%) for up to 8 h. L stands for 1 kb molecular weight ladder. The valency of SC-nbGFP is highlighted in red on top of each lane. M13mp18 is used as a positive control, which shows progressive degradation with increasing FBS concentration and incubation time. Compared to the control group, other SC-nbGFP nanostructures are more stable. However, the reduced intensity of SC-nbGFP bands over time and with increasing FBS concentration indicate some degree of degradation.

**Fig. S7.**
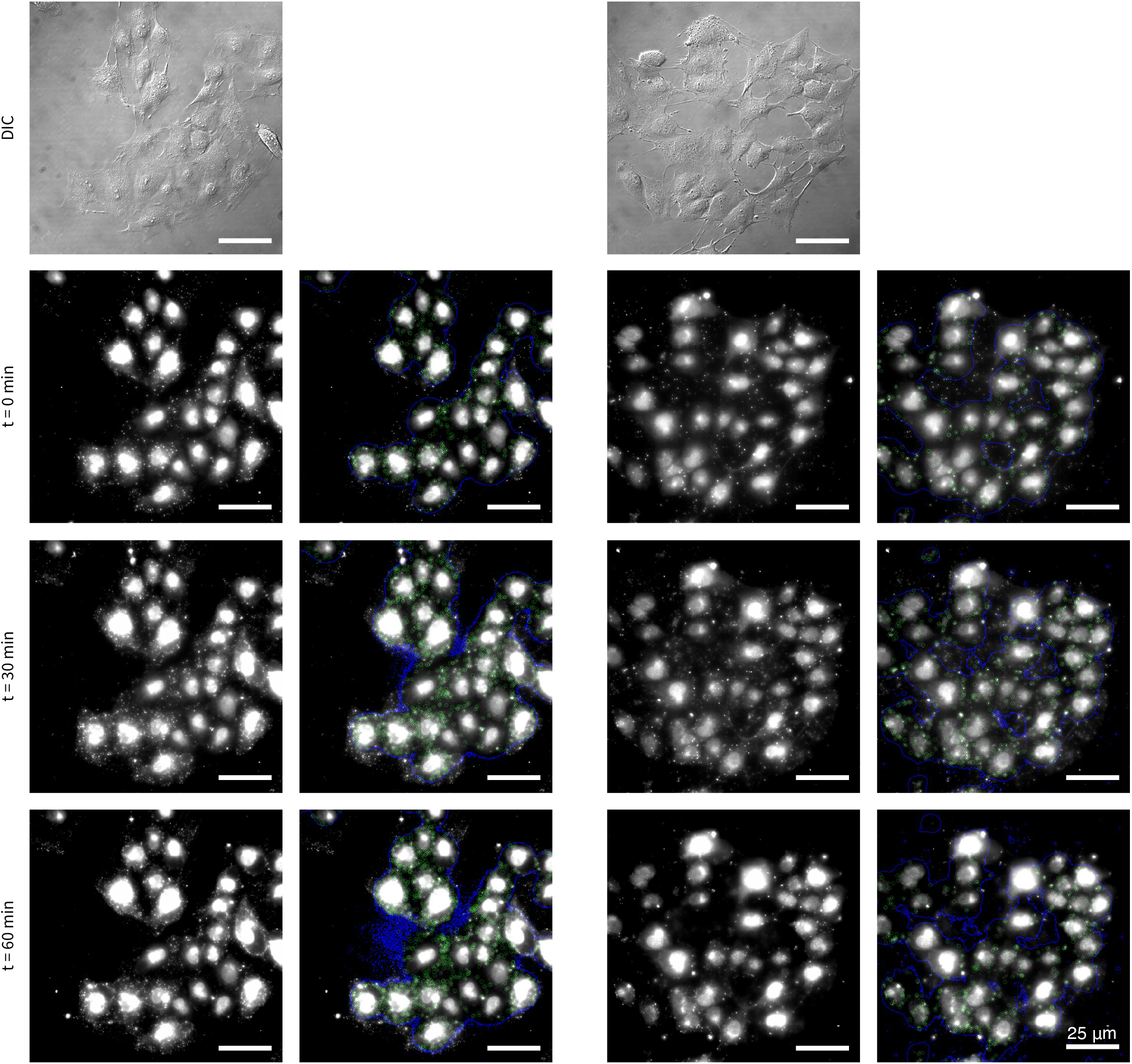
SC-nbGFP blocks viral attachment to host cells at various time points. Representative images from widefield fluorescence microscopy performed on PK15 cells infected with viruses pretreated with either SC60H (left) or SC60H-nbGFP (right). The sec and fourth columns represent annotated images from the first and third columns. Cell nuclei are stained with Hoechst (white). The cell area is marked with blue outlines. Internalized virus capsids are indicated by red circles.

**Fig. S8.**
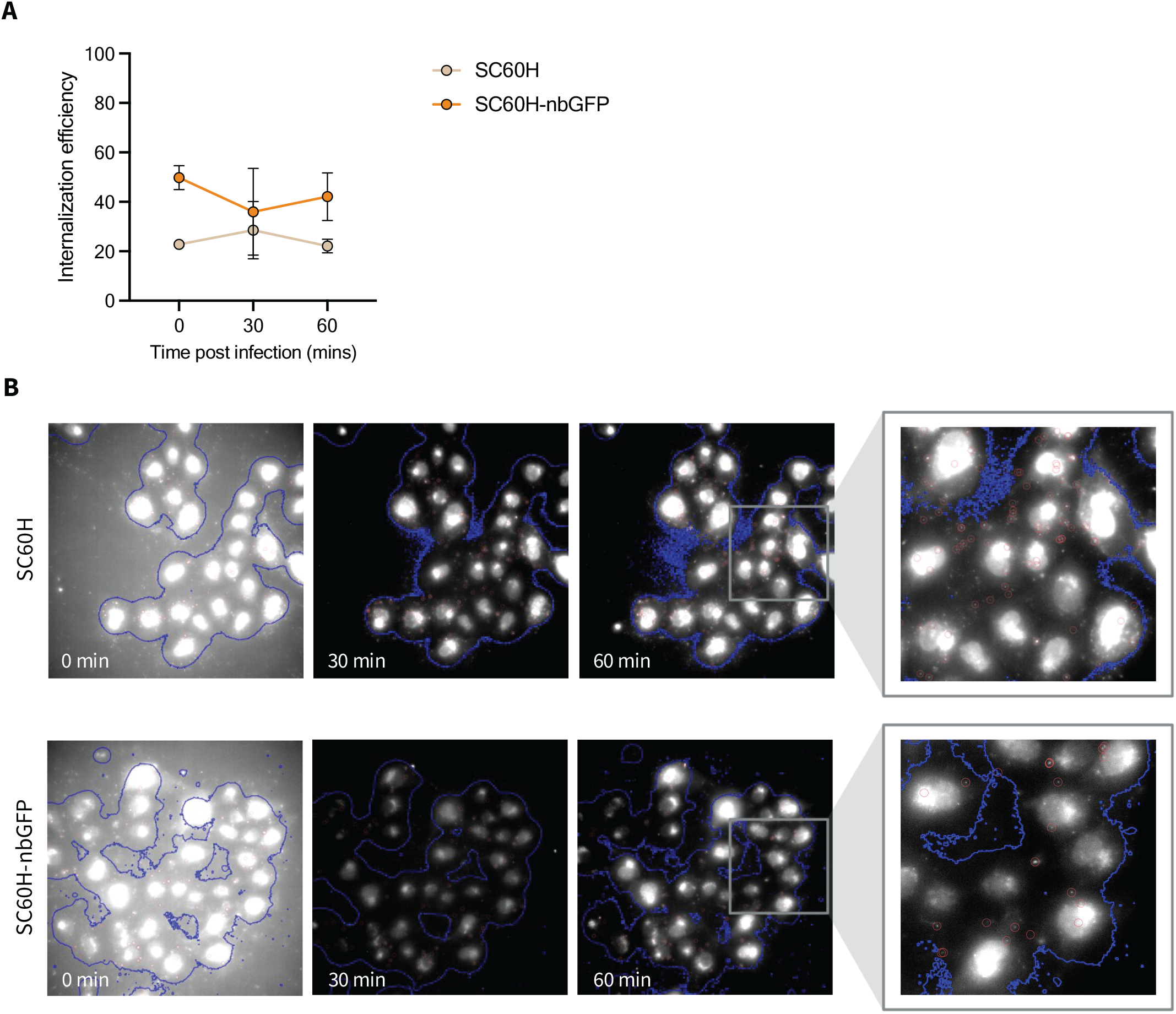
SC-nbGFP does not block viral internalization. (**A**) Internalization efficiency of virus particles pretreated with SC60H-nbGFP versus SC60H (control). Data are presented as mean ± S.D. (**B**) Representative images from widefield fluorescence microscopy performed on PK15 cells infected with viruses pretreated with either SC60H-nbGFP (top row) or SC60H (bottom row). The fourth column represents zoomed-in sections from the third column. Cell nuclei are stained with Hoechst (white). The cell area is marked with blue outlines. Internalized virus capsids are indicated by red circles.

### S2. The IC_50_ is nontoxic to cells in vitro

An ideal drug should have a relatively high therapeutic index (TI), *i*.*e*., it should be effective at low concentrations and toxic only at very high concentrations. Previous studies have shown minimal toxicity of DNA-based molecular devices, making them attractive therapeutic tools. Here, we investigated the cytotoxicity of SC60H-nbGFP in PK15 cells using an LDH assay. Briefly, we incubated cells with different concentrations of SC60H-nbGFP for 24 hours and processed the cell supernatant to quantify cytotoxicity levels. The LDH assay measures the lactate dehydrogenase (LDH) enzyme released by cells upon damage to the plasma membrane in response to cytotoxic components in the cell culture medium.

For the range of concentrations we tested, we observed no apparent cytotoxicity with SC60H-nbGFP at concentrations <10 nM (Figure S4). This is consistent with previous studies employing 2D and 3D DNA origami nanostructures. At concentrations > 10 nM, SC60H-nbGFP exhibited a cytotoxic effect on PK15 cells that was similar to that of the SC60H and M13 DNA controls, indicating that the observed cytotoxicity is due to the DNA component of the multivalent assembly. At the highest DNA concentration of 150 nM, the viability dropped to 80− 90% for SC60H-nbGFP and SC60H. M13’s drop in cell viability was comparatively higher at 70 − 80%. The IC_50_ value of 4.2 ±0.9 nM falls below the toxicity threshold of SC60H-nbGFP. However, additional experiments are needed to understand how this data fits into the broader context of the pharmacological profile of SC60H-nbGFP.

### S3. Aggregation is not the predominant mechanism of virus inhibition by SC-nbGFP

To investigate whether SC-nbGFP nanostructures are aggregating PRV 486 particles, we analyzed the particle size distribution of viruses incubated with multivalent nanostructures using a nanoparticle analyzer (NanoSight NS3000 equipped with Nanoparticle Tracking Analysis (NTA) software; Figure S5). As controls, we incubated viruses with SC (scaffold without binders) and with nbGFP alone (binders without scaffold). For reference, PRV has a diameter of ∼ 200 nm. The NTA data we collected confirmed the size of PRV. In comparison, all the other groups also showed a uniform size distribution of ∼ 200 nm, indicating the absence of the formation of higher-order structures. Our studies of valency found that only one nbGFP per SC was sufficient to produce antiviral effects. This suggests that a) not all binding sites on the virus are saturated, since the size of SC would likely sterically block access from other SC-nbGFP and b) crosslinking interactions between two or more viral particles mediated by the SC are not a critical feature of the inhibition mechanism. These pieces of data provide strong evidence that SC-nbGFP does not function through the aggregation of PRV.

### S4. Serum stability of SCnbGFP nanostructures

The structural integrity of 3D DNA origami nanostructures relies on a minimum concentration (∼ 5 − 20 mM) of cations like Mg^2+^ to stabilize the electrostatic repulsive forces imparted by their negatively charged nucleic acid backbones. 58 We maintained a 12.5 mM concentration of Mg^2+^ throughout the synthesis and purification of SC and SC-associated nanostructures. This concentration is one order of magnitude higher than the physiological concentration found in the human body, which is 0.7-1 mM. ^59,60^ Previous studies have reported the loss of structural integrity of DNA nanostructures in cation-depleted-cell culture media which could affect their functional efficacies. ^29^ Furthermore, serum under physiological conditions can degrade DNA origami nanostructures by the nuclease activity of DNA-degrading enzymes. To characterize the physiological compatibility of SC-nbGFP nanostructures, we evaluated their stability in cell culture media with different FBS concentrations. We incubated SC4H-nbGFP, SC24H-nbGFP, and SC60H-nbGFP nanostructures in 0%, 2%, and 10% FBS media for up to 8 h and assessed them by agarose gel electrophoresis. The M13mp18 DNA was used as a control, which is readily degraded. The presence of distinct bands revealed the maintenance of structural integrity before and after 1 h, 2 h, and 8 h incubation. However, the reduced intensity of the bands with increasing incubation time indicated some degradation (Figure S6).

### S5. Methods

#### Chemicals and kits

Tris-acetate EDTA (TAE) buffer, magnesium chloride hexahydrate (MgCl_2_.6H_2_0), methylene blue, pluronic-F127, casein, dimethyl formamide (DMF), agarose and polyethylene glycol 8000 (PEG8000), triethylammonium acetate (TEAA) were purchased from Sigma Aldrich. Cell culture consumables were purchased from Corning. µ-slide 8-well plates for confocal imaging were purchased from Ibidi. DNA ladders, SYBR gold dye, anti-His HRP, streptavidin-HRP, and TMB substrate were purchased from Thermo Fisher. All ELISA reagents (except for TMB) were purchased from Bethyl Laboratories. The BG-GLA-NHS (#S9151S) was purchased from New England Biolabs. The CytoTox 96 non-radioactive cytotoxicity assay kit (#G1780) was purchased from Promega.

#### Oligonucleotides and DNA templates

A total of 16 DNA origami nanostructures (9 conjugated with nbGFP, 6 conjugated with GFP aptamers, and 1 without handles) were designed using Tiamat.^39^ All DNA staple strands used to assemble scaffolded snub cube (SC) DNA origami nanostructures and modified DNA oligonucleotides for docking virus binders onto the SC scaffold were purchased in 96-well plates from Integrated DNA Technologies at 100 nmol synthesis scale with concentrations normalized to 500 µM. The single-stranded M13mp18 scaffold strand was produced in-house using a previously published protocol. ^61,62^

#### Cell culture and virus propagation

PK15 cells, a transformed porcine kidney cell line, were obtained from ATCC (#CCl-33). Recombinant PRV strains, PRV 486 and PRV 483, were previously described. 13 Cells were maintained in complete Dulbecco’s modified Eagle’s medium (DMEM) supplemented with 10% fetal bovine serum (FBS), 4 mM L-glutamine, 100 U/mL penicillin, and 100 µg/mL streptomycin in a humidified 37 °C, 5% CO_2_ incubator.

Viruses were propagated in PK15 cells as follows: PK15 cells were grown in complete medium to 90% confluency in a sterile 10 cm cell culture dish. After removing the media and washing the cells once with phosphate-buffered saline, they were infected with PRV at a multiplicity of infection (MOI) of 0.01 infectious units per cell, in a final volume of 1 mL. Cells were incubated to allow virus adsorption for 1 h in a humidified 37 °C, 5% CO_2_. After 1 h, the infection inoculum was removed and replaced with 10 mL of fresh virus medium (DMEM supplemented with 2% FBS, 4 mM L-glutamine, 100 U/mL penicillin, and 100 µg/mL streptomycin). Cells were further incubated until 80-90% cytopathic effects were observed, at which point the cells and supernatant were harvested. The mixture was centrifuged at 2000 *×* g for 5 min to remove cell debris, and the virus supernatant was divided into 100 − 200 mL aliquots and stored at − 80 °C. The virus stocks were stored at − 80 °C were thawed in a 37 °C water bath and sonicated in a cup sonicator (10 pulses, one sec on and one sec off for a total of 20 secs at an amplitude of 80%) prior to use.

#### Plaque assay

PK15 cells were seeded in 6-well plates at a density of 4*×* 10^5^cells per well. The next day, cells were washed once with phosphate buffered saline and infected with serial dilutions of virus samples. Cultures were incubated for 1 h in a humidified 37 °C, 5% CO_2_ incubator to allow virus adsorption. The unbound virus was removed and replaced with 3 mL of methocel overlay medium (virus medium supplemented with 2% hydroxypropylmethylcellulose). At 3 days post-infection, the methocel overlay medium was removed and cultures were fixed and stained with a 70% methanol solution containing methylene blue dye and incubated at room temperature for up to 24 hours. The staining solution was removed, rinsed with water, and air dried. To determine the infectious virus titer, the total plaque count was divided by the total volume plated, based on the lowest dilution giving a countable number of plaques, and multiplied by the reciprocal of the corresponding dilution factor.

For the plaque reduction assay, approximately 100 − 200 infectious units of PRV 486 were mixed with different concentrations of SCnbGFP in a final volume of 200 µL DMEM (without supplements) and incubated at 37 °C for 1 hour. A plaque assay was then performed as described above. The residual infectivity (%) was calculated using the control condition as a reference. Inhibition data were fit by nonlinear regression to determine the half-maximal inhibitory concentration dosage (IC_50_).

#### TIRF Fluorescence Microscopy-based detection of interactions between viruses and nbGFP

Corning glass coverslips were used to make flow cells for this experiment. The coverslips were cleaned by sonicating in ethanol, rinsing in ultrapure deionized water, sonicating in acetone, incubating in ethanol, rinsing in ultrapure deionized water, incubating in 2% Hellmanex III cleaning solution, rinsing in ultrapure deionized water, and drying under filtered nitrogen gas. The coverslips were then plasma cleaned for 10 mins (Harrick Plasma; PDC-32G). Immediately after, flow cells were assembled by sandwiching double-sided Kapton tape between a larger and a smaller coverslip. The Kapton tape was cut to include two channels for replicate testing. 10 µL of 5 µM nbGFP was injected into the flow cells and incubated for 10 mins in a humidity chamber. All subsequent wash steps were performed using 200 µL of phosphate-buffered saline. After washing excess nbGFP, 1 mg/mL casein was injected into the flow cells and incubated for 10 mins to block nonspecific binding. The excess casein was washed out, and 10 µL of 100 pM PRV 486 was added to the flow cells. As a positive control, 100 nM EGFP was used. The samples were incubated for 10 minutes and unbound material was washed out. The flow cells were sealed with a coverslip sealant and incubated in a humidity chamber before and in between imaging. Flow chambers were then imaged using an Oxford Nanoimager microscope (ONI) with a 473 nm laser at 2% intensity, or <20 mW, a TIRF angle of 55°, and an exposure of 100 ms.

#### Assembly of snub cube-nanobody GFP nanostructures (SC-nbGFP)

The DNA SC used for nbGFP conjugation was self-assembled in a one-pot reaction in which a 100 nM M13 scaffold was mixed with a 10-fold molar excess of common staple strands, a 10-fold molar excess of the handles corresponding to the valency of the SC, a 10-fold molar excess of the handles that block the remaining spots corresponding to the valency of the SC, and 1 mM TAE + 12.5 mM MgCl_2_. A final reaction volume of 100 µL was annealed in a thermocycler with the following program: 95 °C for 5 mins; 90 °C to 86 °C at a rate of 4 °C per 5 minutes; 85 *°*C to 70 *°*C at a rate of 1 *°*C per 5 minutes; 70 *°*C to 40 *°*C at a rate of 1 *°*C per 15 minutes; 40 *°*C to 25 *°*C at 1 *°*C per 10 minutes; and hold at 4 *°*C at the end of the cycle.

Following annealing, the SC nanostructures were purified from excess staple strands using 100 kDa Amicon spin-column filtration. Columns were passivated for 2 mins with 500 µL of 10% Pluronic-F127 in 1x TAE + 12.5 mM MgCl_2_ and centrifuged at 16000 *×* g for 10 mins. The columns were washed with 500 µL of 1x TAE + 12.5 mM MgCl_2_ before adding the samples and an additional 1x TAE + 12.5 mM MgCl_2_ up to 500 µL. The columns were spun at 1000 *×* g for 15 mins before replenishing the 1x TAE + 12.5 mM MgCl_2_ to 500 µL.

The purified SC scaffolds were mixed with a 10-fold molar excess of BG-conjugated complementary DNA strands and incubated for 90 mins at 37°C. Following annealing, another 2 rounds of 100 kDa Amicon spin-column filtration with passivation were performed following the same procedure described above. Finally, the SC-BG-DNA was mixed with nbGFP at a 5x molar excess of the SC valency in a solution of 1X PBS + 12.5 mM MgCl_2_ + 1 mM DTT and incubated overnight at 4 *°*C with gentle rotation. A final set of 5 rounds of 100 kDa Amicon spin-column filtration with passivation was performed with 1X PBS + 12.5 mM MgCl_2_ used as the wash buffer in place of 1X TAE + 12.5 mM MgCl_2_ and all steps were performed at 4 °C to preserve the stability of the assembled SC-nbGFP. Gel electrophoresis was performed following each purification step to confirm the assembly of the DNA origami nanostructures.

#### Agarose gel electrophoresis

DNA nanostructures were analyzed by agarose gel electrophoresis to assure purity and confirm conjugation. Samples were loaded into a 1% agarose gel according to the following mixture: 1 µL sample, 3 µL ultrapureH_2_O, 1 µL 6X loading dye, and 1 µL 6X SYBR GOLD dye. Along with a 1 kb plus DNA ladder, the samples were run in a buffer of 1X TAE + 12.5 mM MgCl_2_ for 1 hour at 100 V. Gels were imaged with a Bio-Rad Molecular Imager Gel Doc XR System transilluminator at the SYBR GOLD excitation wavelength (495 nm).

#### Synthesis of benzyl guanine conjugated DNA oligonucleotides

3’-amine modified (3AmMO) single-stranded DNA oligonucleotides complementary to the SC overhang handles were diluted in 0.1 M HEPES pH 8.5 to a final concentration of 1 mM. N-hydroxysuccinimide ester-functionalized benzyl guanine (BG-GLA-NHS) was freshly reconstituted in DMF to a 50 mM final concentration. For conjugation, the two solutions were mixed in an oligonucleotide-amine: BG-GLA-NHS molar ratio of 1:10. The final concentration of HEPES was maintained between 50 mM and 100 mM. The reaction was incubated at 4 °C for 16 hours with continuous rotation. After incubation, the reaction was desalted using Bio-Rad micro spin columns and further purified using reverse-phase HPLC to remove unconjugated DNA amine. 100 mM TEAA and 100% methanol were used as the HPLC buffers. The HPLC-purified fractions were lyophilized and reconstituted in water. The concentration of BG-conjugated DNA was determined using a NanoDrop spectrophotometer.

#### Purification of BG-conjugated DNA using reverse phase HPLC

BG-conjugated DNA strands were purified from unreacted DNA using a C-18 column on an Agilent 1220 Infinity LC-HPLC system. Sample DNA mixtures were injected into the column in 50-100 µl volume. The purification was performed using a linear gradient method, with Buffer-A (100 mM TEAA) and Buffer-B (100% methanol). The gradient was run from 10% to 100% of Buffer-B for 40 mins. Migration of DNA and DNA conjugates were monitored using absorbance at 260 nm. Purified volumes of DNA conjugates were collected and further confirmed for their purity and identity by MALDI-TOF mass spectrometry. The purified DNA conjugates were lyophilized and stored at − 20 °C until further use.

#### Mass characterization of BG-DNA conjugates using MALDI-TOF mass spectrometry

All purified products were characterized on an AB SCIEX 4800 MALDI TOF/TOF in the positive ion mode, with 3-Hydropicolinic acid (HPA) as the matrix. Samples were spotted onto a MALDI plate using a sandwich technique (sample-matrix-sample).

#### nbGFP synthesis

The nbGFP protein was expressed from the recombinant plasmid, pBiEX1-nbGFP (AddGene; Plasmid #82711), which was a kind gift from Dr. Sivaraj Sivaramakrishnan (University of Minnesota, Twin Cities, USA). The SNAP-tagged protein construct contained, from the N-to C-terminus: the GFP nanobody (nbGFP), the SNAP tag for oligo labeling, and both FLAG and 6*×* His tags for purification.

The plasmid was transformed into BL21 (DE3) competent E. coli (New England Biolabs), and a single colony of transformed cells was picked from LB-agar plates and used to inoculate a 50 mL culture in LB broth containing carbenicillin (100 µg/mL) antibiotics. This culture was grown for 16 hours at 37 °C, and 250 rpm, at which point it was used to inoculate a 500 mL culture in LB supplemented with carbenicillin at the above concentration. The optical density of the culture was monitored until an OD600 of 0.6-0.8 was reached. It was followed by gene induction using 0.5 mM IPTG for 16 hours at 220 rpm and 18 °C. Cells were harvested via centrifugation at 3000 *×* g for 15 mins at 4 °C. The supernatant was discarded, and the pellet was resuspended in 50 mL of lysis buffer (100 mM NaCl, 25 mM Tris at pH 8.0, 5 mM EDTA 1% Triton-X, 1 mM DTT, and 1X cOmplete protease inhibitor (Roche)) for 1 hour at − 80 °C. After thawing the lysate in RT, it was treated with hen egg white lysozyme (HEWL; Sigma-Aldrich) and DNase I (Sigma-Aldrich), each at a concentration of 1 mg/mL, for 30 mins at 37 °C. The mixture was transferred to an ice bath and sonicated for 10 mins (1 sec on, 2 secs off, 50% amplitude). The lysate was centrifuged at 20000 *×* g for 30 mins at RT to separate cell debris from the periplasmic fraction.

The supernatant containing the nbGFP was loaded directly onto the HisTrap FF (Cytiva) 5 mL column equilibrated with Nickel Wash Buffer containing 25 mM Tris at pH 7.6, 500 mM NaCl, and 10 mM imidazole. To remove nonspecifically bound proteins, the resin was washed with 15 column volumes (CV) of the wash buffer, and the bound proteins were subsequently eluted with buffer containing 25 mM Tris at pH 7.6, 150 mM NaCl, and 500 mM imidazole. The eluted fractions were run on a 15% SDS-PAGE gel to confirm protein expression. The fractions mainly containing pure proteins with 35 kDa bands were pooled together and buffer exchanged using a 3.5 kDa MWCO centrifugal filter unit (Amicon) into an anion exchange buffer containing 20 mM Tris at pH 8.0, 10 mM NaCl. The protein solution was then injected into a HiTrap Q FF anion exchange 5 mL column (Cytiva), equilibrated with the anion exchange buffer, and finally eluted using a buffer containing 20 mM Tris at pH 8.0 and 500 mM NaCl. Nanobody constructs were buffer exchanged into PBS using a 3.5 kDa MWCO centrifugal filter unit, divided into aliquots that were flash-frozen in liquid nitrogen and kept at − 80 °C until further use.

#### ELISA assay

96-well Nunc MaxiSorp flat bottom ELISA plates were coated with 100 µL of PRV 486 at 1 *×* 10^9^ particles/mL, diluted in ELISA coating buffer, and incubated overnight at 4 °C. All the wash steps were performed thrice with 200 µL of 1X Tris Buffered Saline + 0.05% Tween20 (TBST), each wash lasting 5 mins. Wells without virus coating were used as negative controls. Wells with EGFP coating were used as positive controls. After incubation, the plates were washed and blocked with 200 µL of 1 mg/mL casein in TBST for 2 hours at room temperature. Plates were washed, after which 100 µL of nbGFP or nbGFP conjugates or GFP aptamers were added in different concentrations after dilution in TBST + 0.1% BSA and incubated for 1 hour at room temperature. Rabbit anti-pHluorin antibodies were used as positive controls for nbGFP. Plates were washed, and 100 µL of 1:10000 dilution of anti-His Horseradish peroxidase (HRP) for nbGFP, 100 µL of 1:10000 dilution of anti-rabbit HRP for anti-GFP antibodies, and 100 µL of 1:10000 dilution of streptavidin-HRP for GFP aptamers were added and incubated for 1 hour at room temperature. Plates were washed, and 100 µL of 3,3’,5,5’-tetramethylbenzidine (TMB) substrate was added and incubated in the dark for 2− 3 mins at room temperature. The reaction was quenched with 100 µL of 0.15 M H2SO4, and absorbance was read immediately at 450 nm using a microplate reader (Spectra MAX 190, Molecular Devices, Inc.).

#### Flow cytometry-based neutralization assay

PK15 cells were seeded a day before in 24-well cell culture plates at a density of 7.5 *×* 10^4^ cells per well. The cells reached 70− 80% confluency at this seeding density the next day. Approximately 1.5*×* 10^4^ infectious units of PRV 486 were incubated with different concentrations of SC-nbGFP in a final volume of 100 µL for 1 hour at 37 °C. Fresh DMEM was used to dilute the stocks of SC-nbGFP constructs and to constitute the final reaction volume. Before infection, the media from 24-well plates was removed, and the cells were washed once with 1X PBS. The virus mixtures were added to the individual wells, and the cells were incubated to allow virus adsorption for 1 hour in a humidified 37 °C, 5% CO_2_ incubator. After 1 hour, 400 µL of virus medium was added to the wells to make up the final volume of 500 µL per well.

After 48 hours, cells were harvested for flow cytometry. The cell supernatant was removed from the wells and cells were washed once with PBS. Then the cells were fixed with 100 µL of 4% paraformaldehyde (PFA) for 20 mins at room temperature with gentle shaking. Cells were washed and dissociated with 100 µL of 0.15% trypsin for 5 mins at room temperature with gentle shaking. Trypsin was inactivated by adding 200 µL of PBS + 2% FBS. Plates were centrifuged at 300*×* g for 5 mins. The supernatant was discarded, and the cell pellets were reconstituted in 200 µL of PBS + 2% FBS and transferred to individual wells in 96-well flow cytometry round bottom plates. The samples were acquired and analyzed using an Attune NxT Flow Cytometer and software (Thermo Fisher), respectively. In total, 3 *×* 10^4^ single cell events, gated on side scatter area versus height, were recorded for analysis. EGFP was excited with a 488 nm laser, and emission was measured with a 530/30 nm bandpass filter. Untreated cells were used as negative controls and cells treated only with PRV 486 were used as positive controls. The residual infectivity (%) was calculated using the control group with the virus as the reference. Nonlinear regression for dose-response: inhibition was used to curve fit the data and analyze the half-maximal inhibitory concentration dosage (IC_50_).

#### Cytotoxicity assay

Cytotoxicity resulting from treatment with SCnbGFP constructs or individual components was analyzed using the Promega LDH kit. In brief, PK15 cells were seeded in a 96-well cell culture plate a day before at a density of 50000 cells per well. To remove residual LDH activity from the cells, the overnight media was replaced with 100 µL of fresh media. SC60H-nbGFP, SC60H, nbGFP, and M13mp18 constructs were diluted to different concentrations in DMEM and added to the wells at 50 µL per well, for a total of 150 µL. Cells were incubated for 24 hours in a humidified incubator at 37 °C and 5% CO_2_. Untreated cells were used as negative controls. Cells treated with the lysis buffer were used as positive controls and as a reference to calculate the cell viability of other groups. After incubation, the cell supernatant was removed and carefully transferred into individual wells of an optically clear 96-well flat-bottom microplate. A 50 µL LDH reaction mixture was added to the wells, and the plate was incubated in the dark for 30 mins at room temperature. To stop the reaction, 50 µL of the stop solution was added to each well, and the absorbance was read within 1 hour using a microplate reader (Spectra MAX 190, Molecular Devices, Inc.). Cytotoxicity was calculated according to the manufacturer’s protocol, and cell viability was calculated as 1− cytotoxicity.

#### Serum stability assay

The stability of the conjugated snub cube nanostructures was evaluated *in vitro* by incubation at 37 °C for periods of 0, 1, 2, and 8 hours. 20 µL reactions containing 5 nM of the conjugated snub cube and DMEM supplemented with 0, 2, or 10% FBS were incubated for the respective duration before analysis with agarose gel electrophoresis. 2 µL of the sample were combined with 3 µL of water and 1 µL of 6X loading dye and loaded into a 1% agarose gel pre-stained with 1X SYBR GOLD. The samples were run for 1 hour in a running buffer of 1X TAE + 12.5 mM MgCl_2_ at 100 V before visualization with a Bio-Rad Molecular Imager Gel Doc XR System transilluminator at the SYBR GOLD excitation wavelength (495 nm).

#### Particle size distribution analyses

Tracking analysis (NTA) measurements were performed to characterize the particle size distributions of the complexes formed by the interactions between PRV 486 and SC60H-nbGFP. NTA measurements were performed using a NanoSight NS300 instrument (Malvern Panalytical Ltd.), following the manufacturer’s instructions. The virus samples with or without SC60H-nbGFP were serially diluted with PBS to reach a particle concentration of 10^7^-10^9^ particles/mL, suitable for NTA. The samples were injected into the sample unit with 1 mL Luer-Slip sterile syringes (VWR). The capture settings (shutter and gain) and analysis settings were manually set. Each group was run in at least two different sample dilutions, and each sample was analyzed thrice. The video was recorded for 60 secs at 30 fps for each measurement and analyzed using Nanoparticle Tracking Analysis (NTA) 2.0 Analytical software.

To roughly estimate the size of SC60H-nbGFP, we performed a dynamic light scattering (DLS) analysis. DLS measurements were acquired on a Zetasizer instrument (Malvern Panalytical Ltd.). The SC60H-nbGFP samples were diluted in TAE buffer containing 12.5 mM MgCl_2_, to final concentrations of 50 pM, 500 pM, and 5 nM. 1 mL sample volume was loaded into a glass cuvette. Each sample was analyzed twice and the size distributions of samples with an acceptable polydispersity index (PDI) were considered.

#### Widefield microscopy

For widefield fluorescence microscopy experiments, PK15 cells were seeded a day before in a -slide 8-well plate (Ibidi) at a density of 1×10^4^ cells per well. Approximately 10^8^ infectious units of PRV 483 were mixed with 25 nM of SC60H-nbGFP in a final 100 µL volume and incubated at 37 °C for 1 hour. 30 mins before imaging, cells were washed and incubated with Hoechst solution at 1 µM final concentration. The Hoechst solution was removed, and the virus and inhibitor cocktails were added to the cells. Cells were imaged on a Nikon Ti2-E inverted fluorescence microscope using three wavelengths 405 (nucleus), 488 (virus envelope, GFP), and 555 (virus capsid, RFP). Z-stacks consisted of ∼ 10 images per stack, spaced by 0.2 µm, and 4 to 5 fields of view were acquired for each sample. Image analyses were performed using Mathematica.

#### Image analysis

Images were analyzed using a custom Mathematica code that is available at https://github.com/rhariadi/virabloc. Briefly, the blue channel used to image the Hoechst stain was used to manually create a mask of the entire cell boundary. Z-stacks of the red and green channels were manually selected to encompass images containing cell periphery, and those selected images were projected at maximum. The mask was applied to the max projected images. A custom peak finder similar to that previously described was used to identify individual puncta in the green and red channels.63 Colocalization events of the green and red puncta were determined when the centroid distance between individual puncta was within 10 pixels.

#### Statistical analysis

Analyses, including curve fitting, were performed with GraphPad Prism 9. Curve-fitting formulae were stated in the main text. The prism files are available at https://github.com/rhariadi/virabloc/tree/main/Manuscript%20prism%20files. There were no constraints or a priori initial values used in the fitting process in Prism. The *p* values were computed by unpaired two-tailed Student’s t-tests. All measurements are expressed as mean±standard deviation (SD). Significant differences were detected when *p*-value<0.05.

## S6. Supplementary tables

**Table S1:**
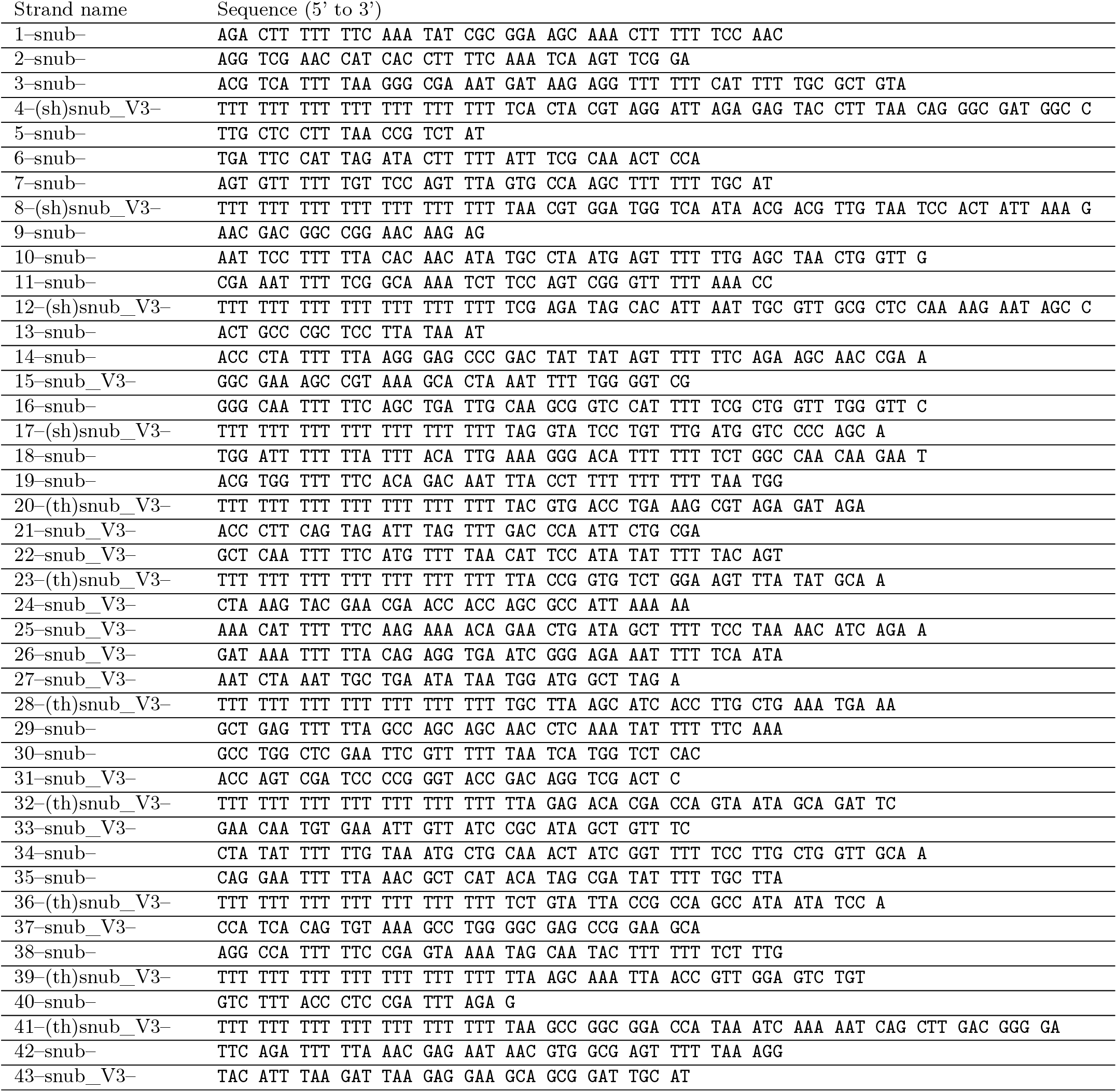
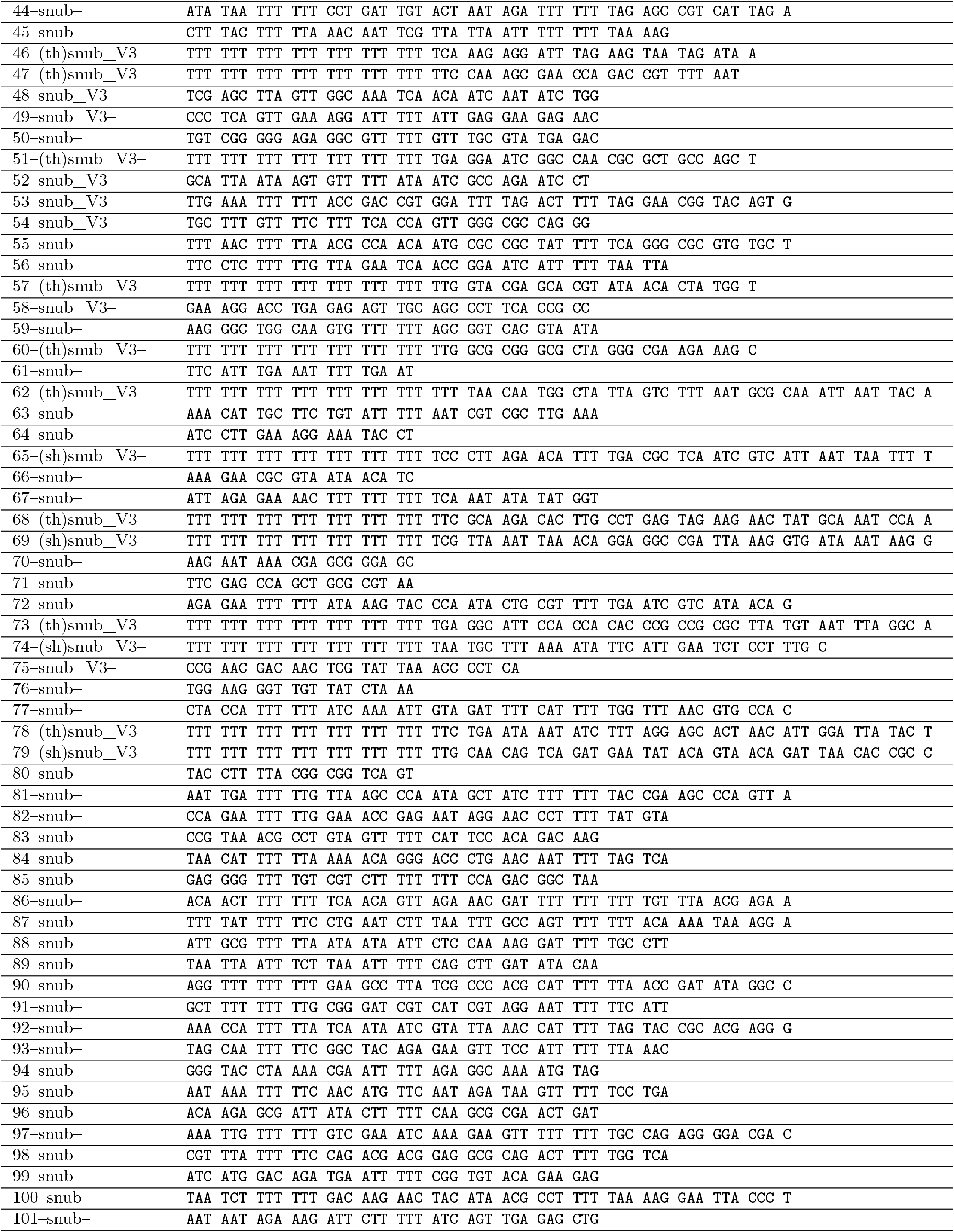
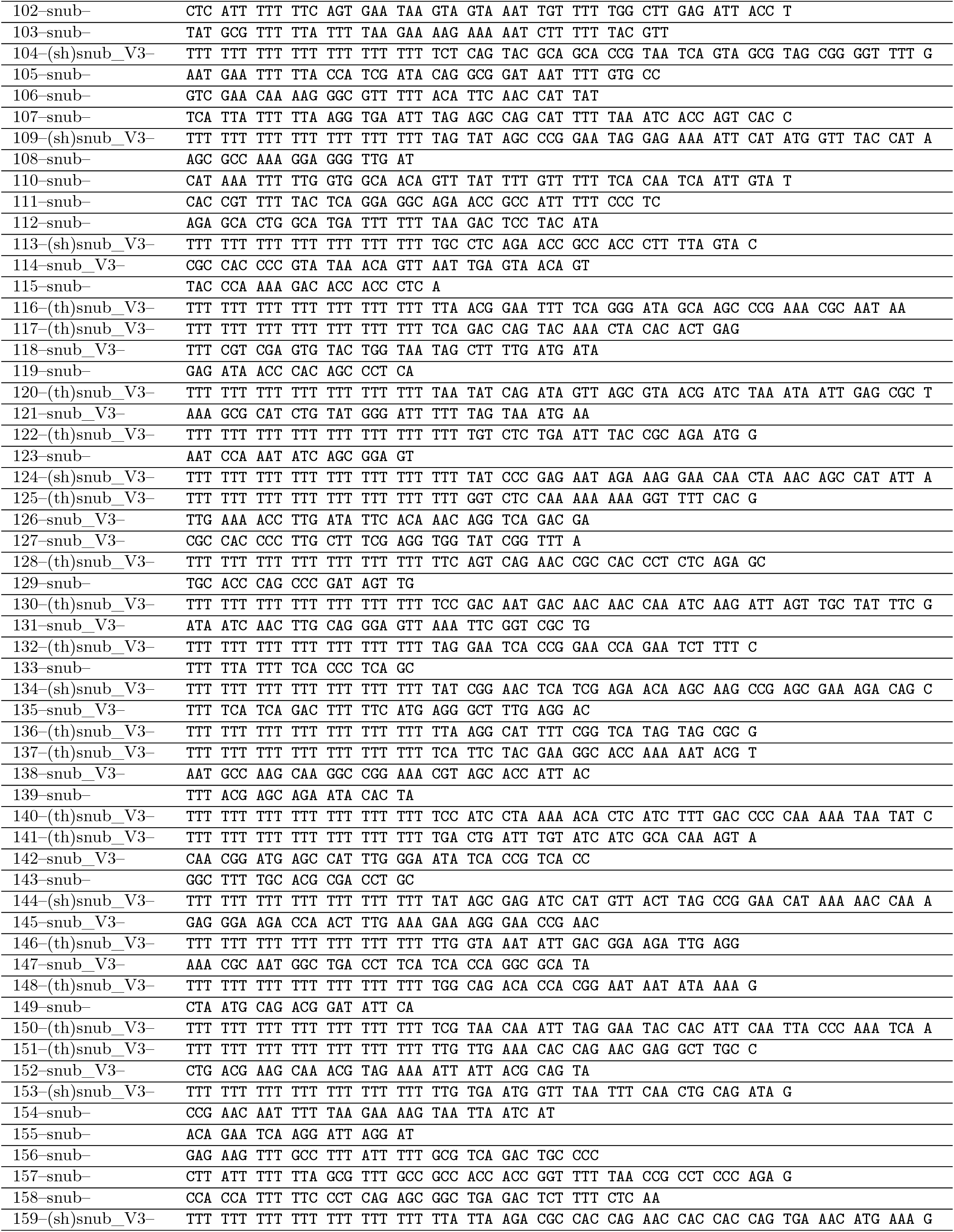
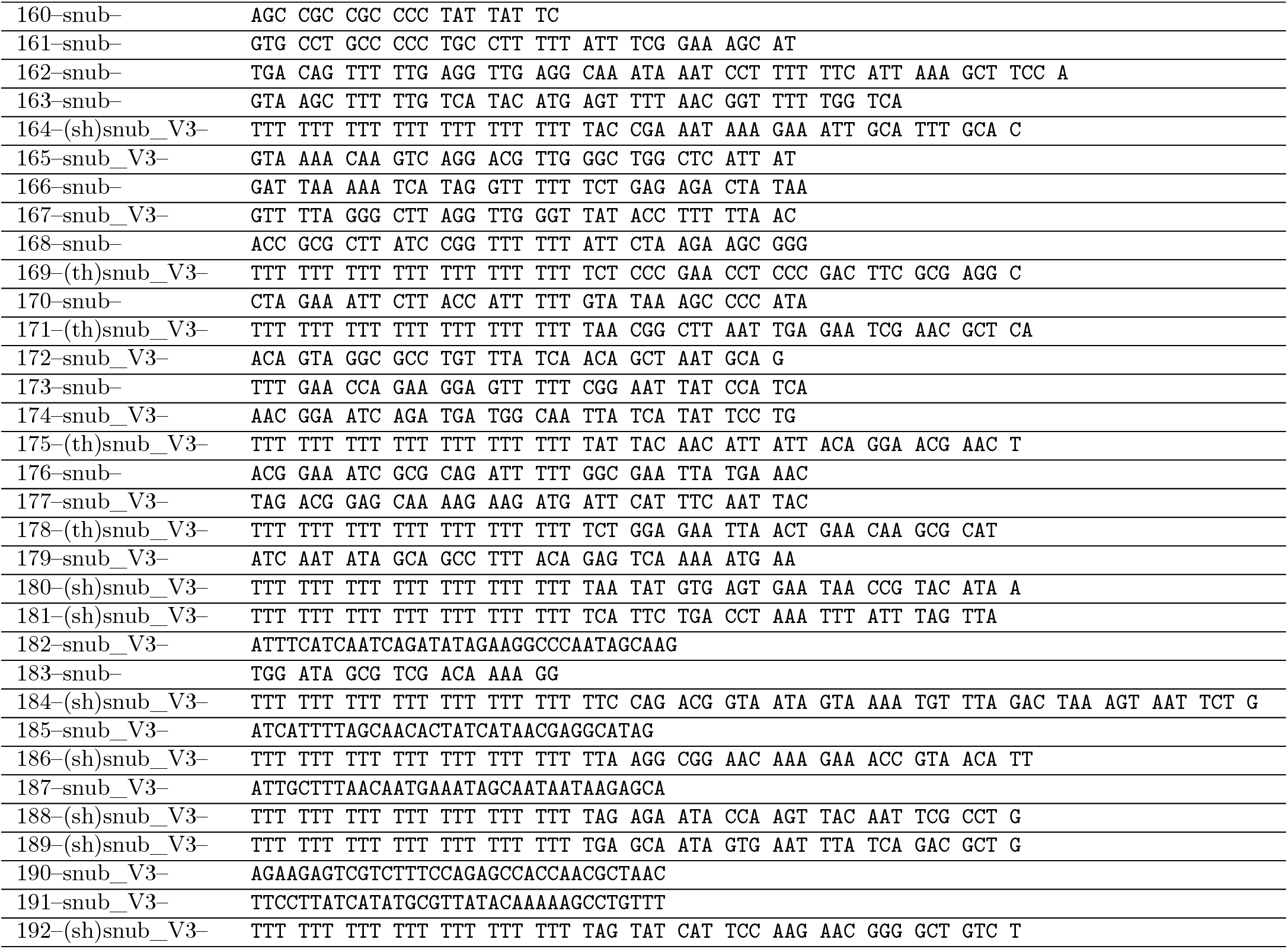

**Table S2:**
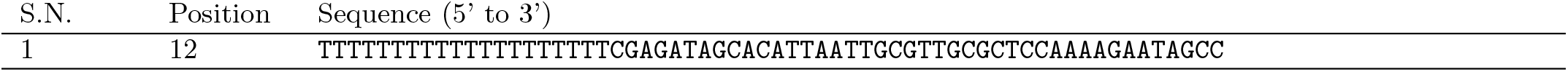
Handle sequence for SC1H.

**Table S3:**
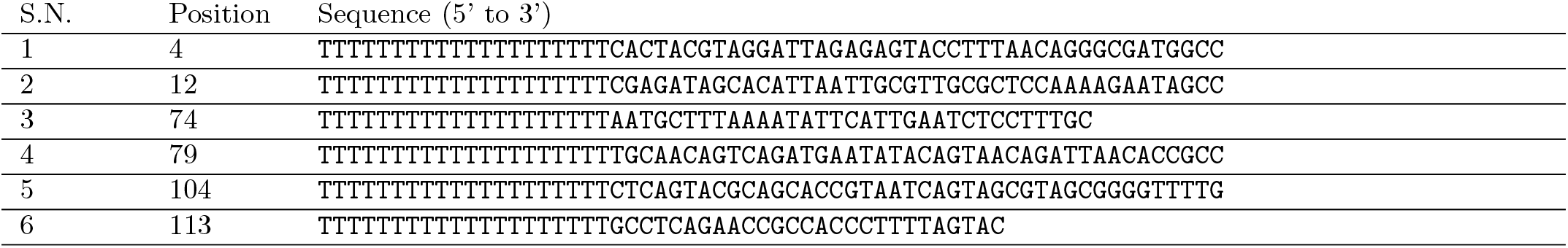
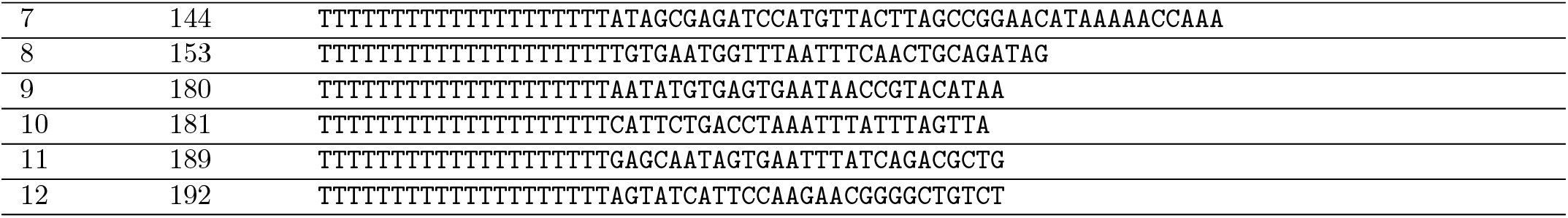
Handle sequences for SC12H.

**Table S4:** Anti-handle sequence.

| Strand name | Sequence (5' to 3') |
| --- | --- |
| Benzyl-guanine-modified Anti-handle | AAAAAAAAAAAAAAAAAAAA-BG |

